# Size-Correlated Polymorphisms in Phyllotaxis-Like Periodic and Symmetric Tentacle Arrangements in Hydrozoan Polyps

**DOI:** 10.1101/2023.08.29.555278

**Authors:** Safiye E. Sarper, Miho S. Kitazawa, Tamami Nakanishi, Koichi Fujimoto

## Abstract

**Introduction:** Periodic organ arrangements occur during growth and development and are widespread in animals and plants. In bilaterian animals, the organs can be interpreted as being periodically arranged along the two-dimensional space and defined by two body axes; on the other hand, in radially symmetrical animals and plants, organs are arranged in the three-dimensional space around the body axis and around plant stems, respectively. The principles of periodic organ arrangement have primarily been investigated in bilaterians; however, studies on this phenomenon in radially symmetrical animals are scarce.

**Methods:** In the present study, we combined live imaging, quantitative analysis, and mathematical modeling to elucidate periodic organ arrangement in a radially symmetrical animal, *Coryne uchidai* (Cnidaria, Hydrozoa).

**Results:** The polyps of *C. uchidai* simultaneously formed multiple tentacles to establish a regularly angled, ring-like arrangement with radial symmetry. Multiple rings periodically appeared throughout the body and mostly maintained symmetry. Furthermore, we observed polymorphisms in symmetry type, including tri-, tetra-, and pentaradial symmetries, as individual variations. Notably, the types of radial symmetry were positively correlated with polyp diameter, with a larger diameter in pentaradial polyps than in tetra- and triradial ones. Our mathematical model suggested the selection of size-correlated radial symmetry based on the activation-inhibition and positional information from the mouth of tentacle initiation.

**Discussion:** Our established quantification methods and mathematical model for tentacle arrangements are applicable to other radially symmetrical animals, and will reveal the widespread association between size-correlated symmetry and periodic arrangement principles.

## 1. Introduction

The periodic arrangement of body parts occurs widely during growth and developmental processes, including segmented structures (e.g., somites in vertebrates) in animals and lateral organs (e.g., leaves and floral organs in angiosperms) in plants. Many animal phyla (bilaterians) share periodic segment arrangements following the bilateral symmetry plane, which is primarily interpreted along two-dimensional (2D) space and defined by the anteroposterior, left–right (L–R), or dorsoventral (D– V) body axes, without an intermediate axis between the L–R and D–V axes to understand the entire picture (Fig. 1, left) (Dequéant and Pourquié, 2008). In contrast, in radially symmetrical animals with multiple symmetry planes, such as cnidarians, organs are periodically arranged in a three-dimensional (3D) space, rather than a 2D space, around the oral–aboral (O–A) axis (Fig. 1, middle), similar to organ arrangements around plant stems (Fig. 1, right) (Bhatia and Heisler, 2018). Unlike studies on bilaterians and plants, quantitative studies on periodic organ arrangements in radially symmetric animals are scarce, and how to establish radial symmetry in animals remains unclear.

**Figure 1.**
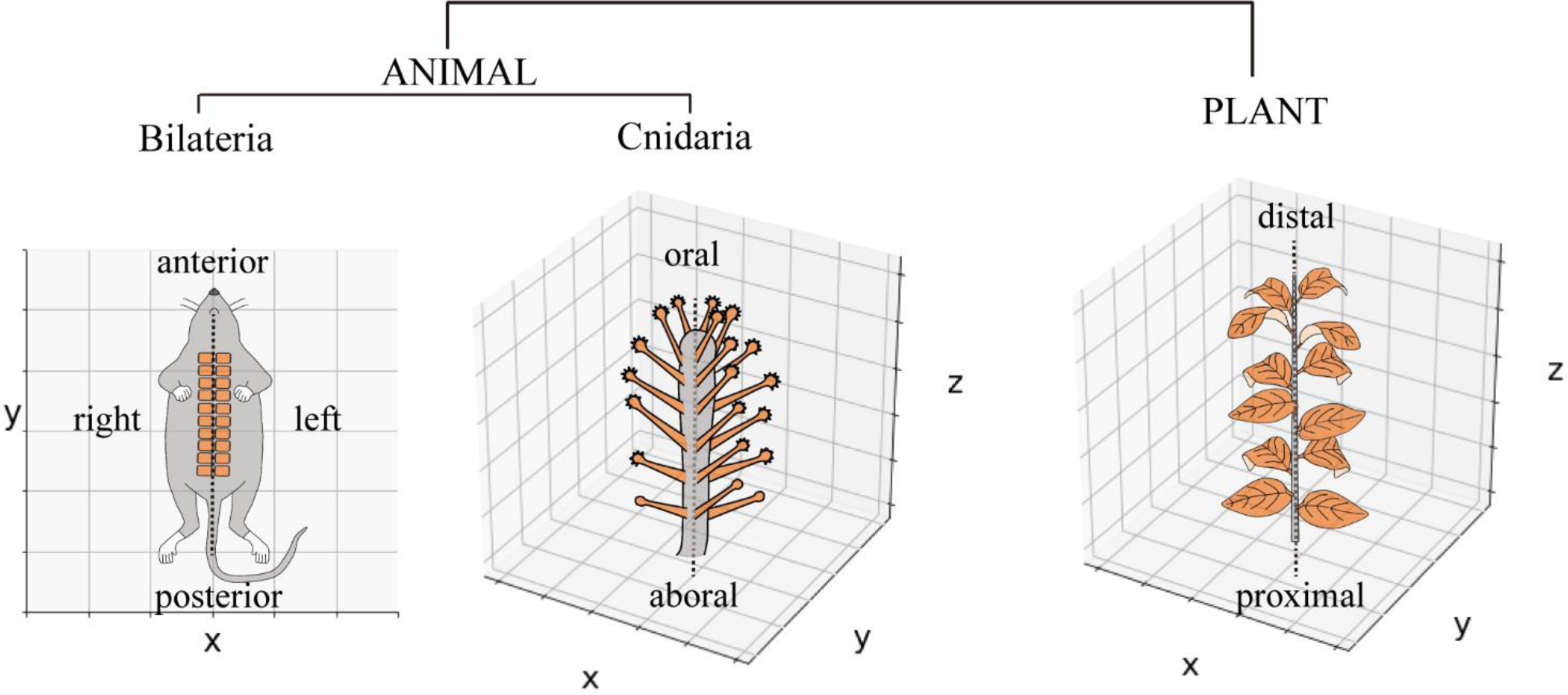
Periodic organ arrangements in animals and plants. Schematic diagram of somite arrangement in a mouse via the single-body axis represented in a two-dimensional space. The tentacle arrangements of *Coryne uchidai* (Cnidarian, Hydrozoa) around the oral–aboral axis and leaf arrangements around the plant stem are represented in a three-dimensional space.

Plant phyllotaxis is a well-established system for quantitatively evaluating periodic organ arrangements with radial symmetry. Recent studies using live imaging have revealed the developmental process involved in the periodic arrangements of organ initiation (Heisler et al., 2005). The quantitative analyses of the distance and angle between organs over 100 years have revealed that phyllotactic patterns are largely classified into two types: whorled (concentric), in which organs are arranged around a plant stem at the same level (distance from the apex), thereby forming a whorl, and spiral patterns, in which organs are individually arranged at different levels at constant longitudinal and angular intervals (Fig. 3A, B) (Green et al., 1998, Bursill and Rouse, 1998). Moreover, quantitative evaluation and classification of periodic organ arrangements have revealed intraspecific polymorphisms in the organ number and arrangement of flowers (Kitazawa and Fujimoto, 2020). Therefore, the whorled or spiral organ arrangement in 3D space and polymorphisms are candidates for the common spatial arrangement principles that can be quantitatively examined between plants and animals with radially symmetric periodicity.

Are there similar spatial arrangement principles and polymorphisms among radially symmetrical animals? Hydrozoan polyps have a simple cylindrical body with a mouth in the oral area and are surrounded by multiple tentacle organs, establishing the radial symmetry of the organism (Fig. 1, middle). *Coryne uchidai* (Cnidaria, Hydrozoa, Corynidae), a hydrozoan polyp, is a potentially suitable, yet uninvestigated, model for quantitatively analyzing organ arrangements in 3D space, where multiple tentacles are formed around the O–A axis, similar to plant organs (Fig. 1A, middle and Fig. 2C) (Hirai and Kakinuma, 1960). Moreover, some hydrozoans exhibit polymorphisms in tentacle numbers, displaying different radial symmetries (Beklemishev, 1969); nevertheless, tentacle arrangements have not been quantitatively analyzed. To this end, we examined the tentacle arrangements of *C. uchidai* to reveal the development of radial symmetry and the emergence of polymorphisms in radial symmetry.

**Figure 2.**
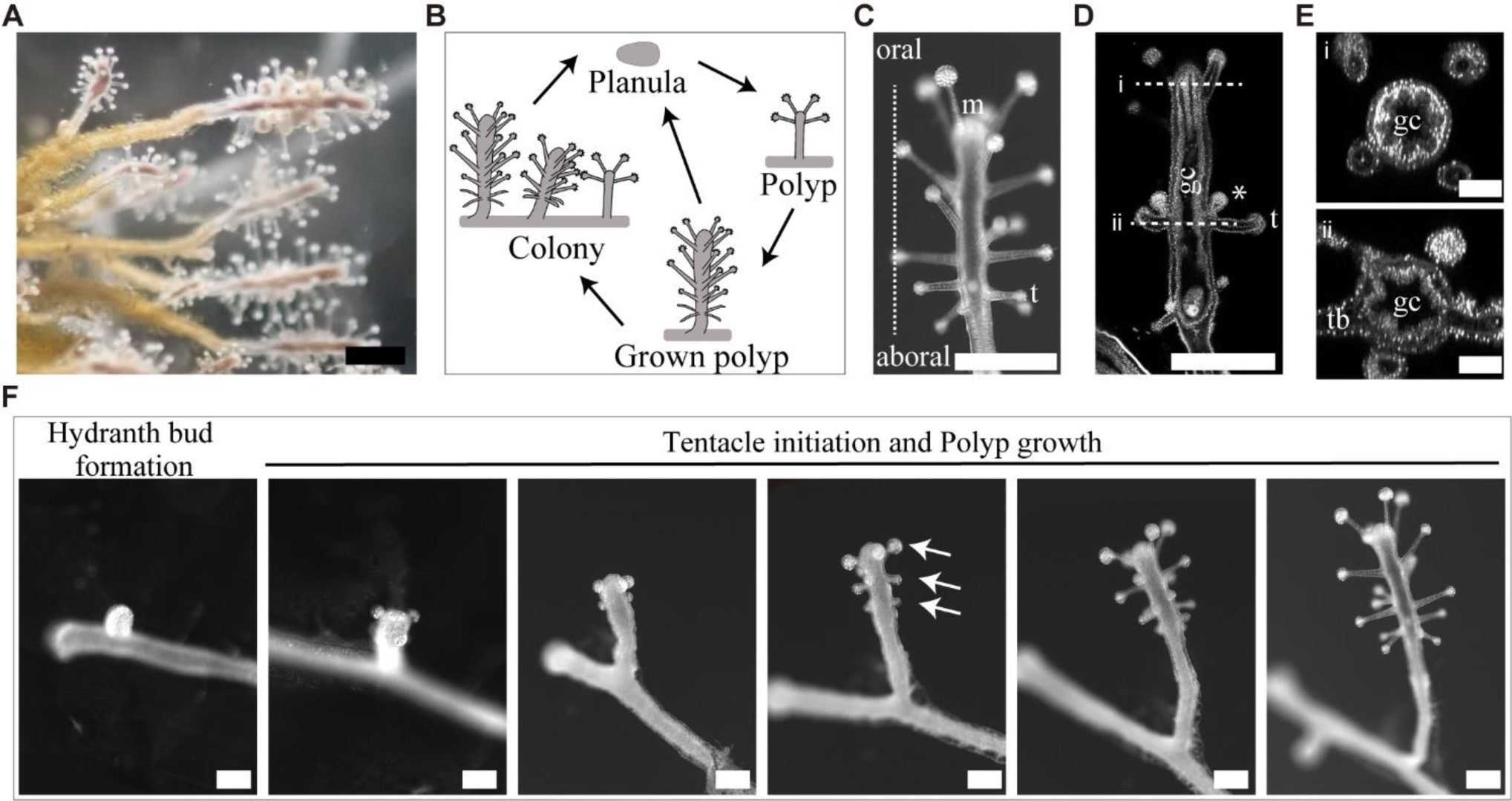
Morphology, life cycle stages, and tentacle initiations of *Coryne uchidai*. **A** External view of a colony of *C. uchidai*. Scale bar: 500 μm. **B** Schematic view of the life cycles of sexual and asexual reproductive modes. Sexual reproduction includes the planula stage. Hydroid polyps can asexually reproduce by extending tube-like stolons, followed by bud formation, which develops into hydroid polyps and colonies. **C** External view of the grown polyp. Scale bar: 500 μm. m: mouth, t: tentacle. **D** Longitudinal section representing the gastrovascular cavity (gc), tentacle (t) arrangements, and gonophores. Asterisks indicate gonophores. Scale bar: 500 μm. **E** Horizontal sections of a polyp demonstrating the gastrovascular cavity (i) and tentacle bases (tb in ii). Scale bar: 100 μm. **F** Live imaging of the stages of asexual reproduction, including hydranth bud formation, transformation, tentacle initiation, and polyp growth. Arrows indicate the ring levels. Scale bar: 250 μm.

## 2. Materials and Methods

### 2.1 Sample collection and nursery

All *C. uchidai* samples were collected between April and June 2022 from Akashi, Hyogo Prefecture, Japan. Forceps were used to gently detach the polyp colony from the substrate. In the laboratory, polyp samples were stored in artificial seawater at room temperature (23 °C–27 °C) with a 12-h day/night cycle and fed *Artemia salina* (brine shrimp) once a week throughout the experimental process. All animal experiments were approved by the Institutional Animal Care and Use Committee of RIKEN, Kobe Branch.

### 2.2 Genomic DNA extraction and species determination

Genomic DNA was extracted from the polyps using the QIAamp DNA Micro Kit (QIAGEN, Germany). PCR amplification of ribosomal DNA (28S) and cytochrome oxidase subunit I (COI) was performed. Mitochondrial COI and nuclear 28S sequences of the collected polyps were amplified via PCR using the following forward and reverse primers: COI, F (5′- GTACTTGATATTTGGTGCTTTTGCAGGCATGGT-3′) and R (5′- CCTAGAAAAGCTATAGCTAATTGAGCGTATACC-3′), and 28S, F (5′’- GCTTAAAATCTCTGTTGCTTGCAACAGCG-3′) and R (5′- CAAGCAAGTGCAAATGCCAATTGTCTG-3′) (Nawrocki et al., 2010). The amplified samples were sequenced by Azenta Life Sciences (Chelmsford, USA). These polyps were confirmed as *C. uchidai* based on their 28S ribosomal and COI DNA sequences.

### 2.3 Clearing, 4′,6-diamidino-2-phenylindole (DAPI) staining, and confocal imaging

To analyze the tentacle organs relative to the polyp body, the polyp samples were fixed with 4% paraformaldehyde after treating them with MgCl_2_ in nursing water. The fixed samples were stained with DAPI (Roche, Basel, Switzerland) and cleared using the CUBIC clearing method (Omnipaque 350, GE Healthcare Japan) to observe their morphology. A confocal fluorescence microscope (Leica SP8, Wetzlar, Germany) was used for imaging.

### 2.4 Quantitative analysis of tentacle arrangements

We employed quantitative analysis methods of plant phyllotaxis to analyze tentacle patterns. The design principles of periodic organ arrangements in plants are categorized into two types: whorls and spirals. These two categories were distinguished by measuring internode lengths (Δ*h*) and divergence angles (φ) (Fig. 3A-M), namely, the distances of two successive organs along the stem (Δ*h*_*i*_ = *h*_*i*+1_ − *h*_*i*_) and projection angles of two successive organs (φ_*i*_ = *θ*_*i*+1_ − *θ*_*i*_, where *θ*_*i*_ denotes the angular position of the *i*-th organ) relative to the stem. In the whorled arrangement, organs are arranged in multiple whorls around the stem; therefore, the internode lengths are approximately zero in each whorl, with a large gap between whorls; on the other hand, in the spiral arrangement, both the internode lengths and divergence angles are not zero but constant throughout the stem (Fig. 3D, E, K, L). The whorled arrangement is further characterized based on the number of organs present in a whorl. The phyllotaxis measurement system was used to perform measurements in hydrozoan polyps. Because hydrozoan polyp bodies demonstrate considerable curves and thickness, mouth position was used as a reference, and the longitudinal distances between the mouth and each tentacle along the O– A axis were measured (where *h*_*i*_ denotes the position of the *i*-th organ referenced to the mouth position) and the internode length of two successive tentacles was calculated (Fig. 3C, F). Tentacle indices (*i*) were ordered by the distance from the mouth and arranged in ascending order. To quantify angular arrangement, an angular coordinate was used for each tentacle base position with the coordinate origin at the polyp center and a certain position of the 0° polyp (hereafter referred to as the position angle, *θ*) (Fig. 3J) and the difference in the angle between the tentacles and the neighboring angular positions within a ring was measured (Fig. 3R). To evaluate the regularity of the angular positions, the circular mean 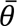 (Equation 1), the resultant vector *R_v_* (Equation 2), and circular standard deviation *S* (Equation 3) of the differences in position angles were measured using the following functions in Python 3:

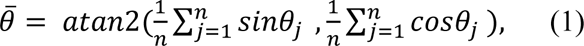

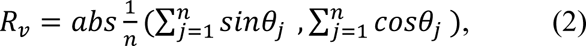

and

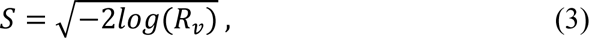

where *j* denotes the tentacle indices ordered by the position angle and arranged in ascending order (Mardia and Jupp, 2009). Here, we demonstrated the circular mean angle and circular standard deviations as; 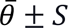.

**Figure 3.**
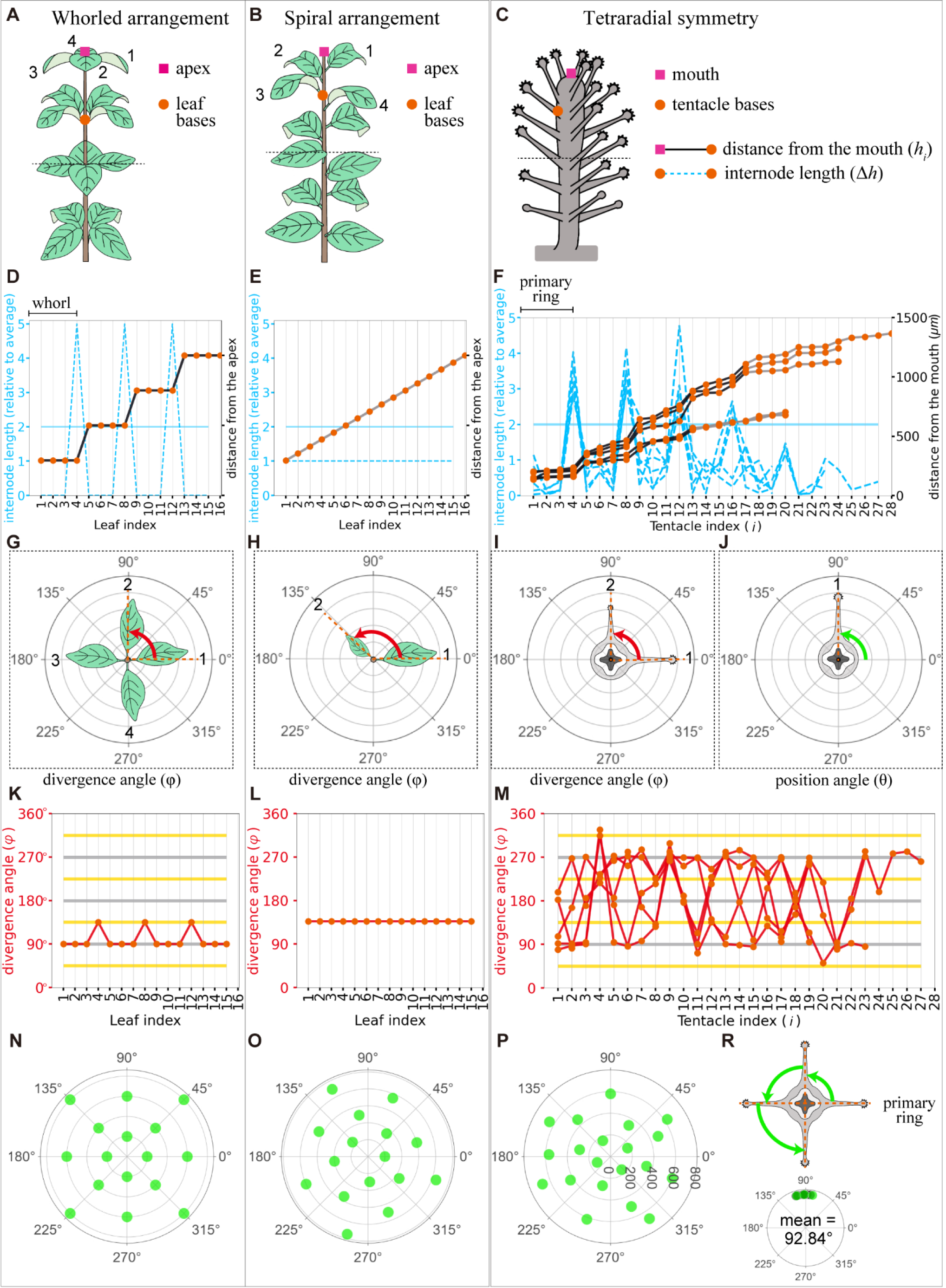
Plant phyllotaxis and *Coryne uchidai* tentacle arrangement measurements. **A-C** Schematic view of whorled leaf arrangement **(A)**, spiral leaf arrangement **(B)**, and *C. uchidai* tentacle arrangement **(C)**. The apex of the plants/mouth of the polyp (pink squares) and leaf/tentacle organ bases (orange circles) are shown. **D-F** Internode length (blue dashed lines) of the successive organs and distance from the mouth (black or grey line) as a function of organ indices in the whorled leaf arrangement **(D)**, spiral leaf arrangement **(E)**, and *C. uchidai* tentacle arrangement (n = 5) **(F).** Black plot lines connecting the orange circles indicate the presence of the ring or whorled arrangement, whereas gray plot lines indicate their absence. **G-J** Schematic views of the divergence and position angles evident in the horizontal slices. Horizontal slices framed with dotted lines reveal the divergence angles (red) of successive leaves **(G-H)** and tentacle organs **(I)**. Measurement of the position angle (green) of the angular coordinate of each tentacle with the coordinate origin at the polyp center **(J). K-M** Divergence angles (red lines) of the successive organs indicated by orange circles as a function of organ indices in the whorled leaf arrangement **(K)**, spiral leaf arrangement **(L)**, and *C. uchidai* tentacle arrangement (n = 5). The measured divergence angles of the tentacles did not reveal a clear pattern **(M)**. The grey and yellow horizontal lines indicate multiples of period angles (120° in tri-, 90° in tetra-, and 72° in pentaradial symmetries) and their half, respectively. **N-P** Polar plots show the angular positions of the organs as a function of the distance from the apex/mouth in the whorled leaf arrangement **(N)**, spiral leaf arrangement **(O)**, and tentacle arrangement **(P)**. Angles between the nearest tentacles within the primary ring in the representative sample and other samples indicated with green and black dots, respectively **(R).** Datasets are identical for **F, M, and R.**

### 2.5 Building a mathematical model

We built a model for tentacle arrangement by combining the present measurements with previous models for *Hydra* organ arrangements (Meinhardt, 2012). We hypothesized regulations by two inhibitory morphogens and one activatory morphogen diffusing on the polyp body surface (Fig. 6A), which were represented as cylindrically arranged cell populations. The following reaction–diffusion equations represent the spatiotemporal kinetics of the morphogens:

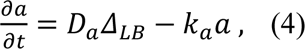

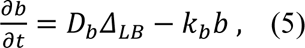

and

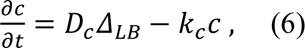

where *a* denotes the concentration of the activator A; *b* and *c* denote the concentrations of inhibitors B and C, respectively; *D_a_*, *D_b_*, and *D_c_* denote the diffusion coefficients; Δ_*LB*_ denote the Laplace-Beltrami operator calculating the diffusion on cylindrical polyp surface; and *k_a_*, *k_b_*, and *k_c_*denote the degradation rates. Both A and B were synthesized in the mouth area at constant rates of *s_a_* and *s_b_*, respectively; on the other hand, C was synthesized in the tentacles at a constant rate of *s_c_*(Fig. 6B). Supplementary Table S1 presents the parameter values for these equations. Numerical simulations of the model were performed using the Euler method, a finite difference scheme, on the Python-based CompuCell3D platform (Swat et al., 2012) under Neumann boundary conditions.

## 3. Results

### 3.1 Tentacle initiations in *C. uchidai*

Tentacles were spatially periodically arranged in *C. uchidai*. Multiple tentacles formed ring-like arrangements that were repetitively positioned in the 3D space along the O–A axis (Fig. 2C-E). We also performed time-lapse imaging of the growing polyps of *C. uchidai* to reveal where and how tentacle initiations proceed (Fig. 2F). Following 5–18 hours after the growth of polyp stolons and the transformation of hydranth buds into hydranths, multiple tentacles were initiated in a major fraction of the polyps (n = 5; Fig. 2F, two panels from the left end), almost simultaneously at the oral side of the polyp, forming a ring-like arrangement (Fig. 2F, third panel from the left end). The other rings formed toward the aboral side based on the subsequent initiations of tentacles as the polyp body grew longitudinally (Fig. 2F, third panel from the left end). Therefore, live imaging revealed a developmental time course in which the initiation of multiple tentacles within each ring was simultaneous, while rings initiated sequentially from the oral to the aboral side, thereby forming a periodic arrangement during the growth process of polyps.

### 3.2 Quantitative analyses of tentacle arrangement

Next, we examined whether periodic ring-like arrangements are commonly observed in other polyps and whether and what radial symmetry is observed in the rings. To quantitatively analyze the spatial arrangement of the tentacles, we used a plant phyllotaxis measurement system (Fig. 3A-J). By comparing the internode lengths between each tentacle, a ring-defining gap was defined; this gap could distinguish the different rings observed during live imaging (Fig. 2F, fourth panel from the left end). The internode length was two times larger than the average of all internode lengths of the sample 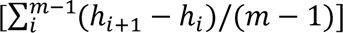, where *m* denotes tentacle number (Fig. 3F). Analysis of five polyps that form approximately 20–28 tentacles revealed ring-defining gaps after the first four tentacles around the mouth corresponding to the ring (hereinafter referred to as the primary ring) (n = 5) (Fig. 3F). Ring-defining gaps appeared in every four tentacles at the first several tentacles from the mouth and resembled the whorled arrangement of plant phyllotaxis (Fig. 3D, F). This whorled arrangement was evident in regularly spaced organs within the whorl (Fig. 3G, N). Thus, we examined the angle between the nearest tentacles within the primary ring (Fig. 3J, R). The mean angle was 92.84°±9.4° (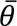 ± *S*, n = 5), which was close to 90°, indicating tetraradial symmetry (Fig. 3R).

### 3.3 Polymorphisms in radial symmetry

While the major polyp samples exhibited tetraradial symmetry, we observed considerable differences in the tentacle numbers and position angles of the other samples. First, we counted the number of tentacles in the primary ring based on ring-defining gaps. Most polyps exhibited four tentacles in the primary rings (41 polyps), followed by three (13 polyps; Fig. 4A, upper panel), five (12 polyps; Fig. 4B, upper panel), six (4), nine (2), and 10 tentacles (1). In polyps with three tentacles in the primary ring, the mean angle between the nearest tentacles was ∼120° (119.45°±14.18°), indicating triradial symmetry (Fig. 4A, lower panel). Similarly, in samples with five tentacles, the mean angle between the nearest tentacles was ∼72° (72.72°±8.93°), indicating a pentaradial symmetry (Fig. 4B, lower panel). Therefore, using tentacle numbers within the primary ring, we could define three types of radial symmetries, with the most frequent type being tetraradial symmetry (56.16%), followed by tri- (17.8%) and pentaradial (16.43%) symmetries.

**Figure 4.**
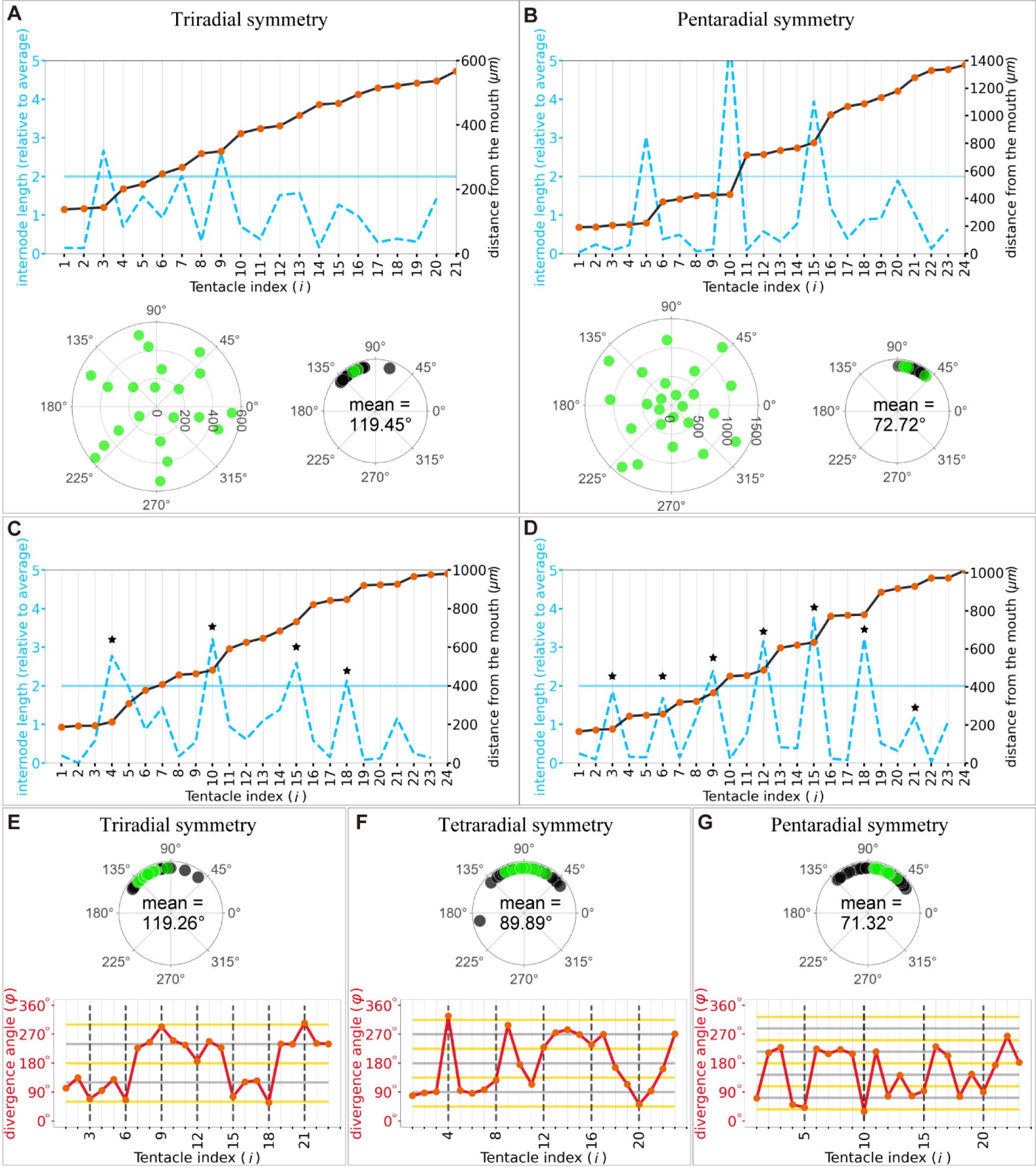
Periodicity of the tentacle arrangement and regularity with polymorphic symmetries. **A-B** Internode length (blue dashed lines) of the successive tentacles and distance from the mouth (black plot lines) as a function of tentacle indices in tri- **(A)** and pentaradial **(B)** symmetries identified in the primary ring **(upper panel)**. Polar plots of the tentacles **(lower left panel)** and angles between the nearest tentacles within the primary ring in the representative sample and other samples indicated with green and black dots, respectively **(lower right panel)**. **C** Samples demonstrating different numbers in the successive rings evident in the ring-defining gap criteria. Star marks indicate each period. **D** Samples exhibited periodicity in every three tentacles. Star marks indicate each period. **E-G** Angles are measured based on the differences in the position angles in tri- **(E, upper panel)**, tetra- **(F, upper panel)**, and pentaradial **(G, upper panel)** symmetries in the whole body in the representative sample and other samples indicated with green and black dots, respectively. Divergence angles (red lines) of the successive tentacles as a function of tentacle indices in tri- **(E, lower panel)**, tetra- **(F, lower panel),** and pentaradial **(G, lower panel)** symmetries. The grey and yellow horizontal lines indicate multiples of period angles (120° in tri-, 90° in tetra-, and 72° in pentaradial symmetries) and their half, respectively. Black dashed vertical lines indicate the numbers for each period.

To determine whether the symmetry type in the primary ring was present in other rings in the whole body, we quantitatively analyzed and classified the periodicity of the arrangements. In most samples (36 of 40 polyp samples with ≥20 tentacles), the ring-defining gap providing ring arrangements appeared more than three times in each polyp, indicating the periodicity of the arrangements. Among them, the majority displayed the same number of tentacles in successive rings (Fig. 4B); however, some polyps did not display the same number of tentacles (e.g., 4, 6, 5 tentacles; Fig. 4C). To quantify the periodicity in the tentacle arrangements in the whole body, each internode length was measured and compared to determine the period in which a significant internode length appeared in each sample. The correlation coefficient R*_k_* between successive internode lengths *h*_*i*+1_ − *h*_*i*_ and *h*_*i*+1+*k*_ − *h*_*i*+*k*_(*k* = 1,2,……, 8) for each polyp with ≥20 tentacles was calculated (n = 40). When large gaps periodically appeared in every *x* tentacles, correlation R*_k_* became the largest at *k* = *x*, e.g., R_5_ = 0.79, whereas |R*_k_*| < 0.32 (*k* ≠ 5; Fig. 4B). Therefore, *k* that provides the maximum value of R*_k_* was regarded as an indicator of the periodicity of cycle *k* and was used to classify the periodicity based on *k* when max{R*_k_*} > 0.5 (Table 1). As a result, most samples with tetra- and pentaradial symmetries in the primary ring exhibited periodicity in every four and in every five tentacles, respectively, throughout the body. Furthermore, samples with six and nine tentacles in the primary ring demonstrated periodicity in every three tentacles (Fig. 4D). Therefore, polymorphisms observed in the primary ring carried on the whole body during periods three, four, and five.

**Table 1.** Classification of symmetry types based on the primary ring-defining gap and ring periodicity defined with correlation coefficients.

|  |  | Periodicity of tentacle pattern of the whole body<sup>1</sup> |  |  |  |  | Total |
| --- | --- | --- | --- | --- | --- | --- | --- |
|  |  | 3 | 4 | 5 | 6 | n.s.<sup>2</sup> |  |
| Primary<br>ring<br>based on<br>ring-defining<br>gap | 3 | 0 | 0 | 0 | 0 | 1 | 1 |
|  | 4 | 0 | 11 | 2 | 0 | 10 | 23 |
|  | 5 | 0 | 0 | 7 | 1 | 2 | 10 |
|  | 6 | 1 | 0 | 0 | 1 | 2 | 4 |
|  | 9 | 1 | 0 | 0 | 0 | 0 | 1 |
|  | 10 | 0 | 0 | 0 | 0 | 1 | 1 |
| Total |  | 2 | 11 | 9 | 2 | 16 | 40 |

<sup>1</sup> $k$ of max $\{R_k\}$
<sup>2</sup> $p > 0.05$

Next, to evaluate the symmetry of the tentacle arrangements in the entire body, regularity within the rings was determined by measuring the position and divergence angles. In samples exhibiting periodicity in every three, four, and five tentacles, the mean angles between the nearest tentacles within the same ring were approximately 120° (119.26°±16.03°, n = 15), 90° (89.89°±14.92°, n = 16), and 72° (71.32°±15.32°, n = 12), indicating tri-, tetra-, and pentaradial symmetries, respectively (Fig. 4E, F, G, upper panels). In addition, the divergence angles between successive rings were half of these angles, i.e., approximately 60° (or 180° and 300°), 45° (or 135°, 225°, and 315°), and 36° (or 108°, 180°, 252°, and 324°) in the tri-, tetra-, and pentaradial samples, respectively (Fig. 4E, F, G, lower panels). Half of the mean angle between the nearest tentacles indicated alternate arrangements in successive rings, which are common in whorled plants (Fig. 3K). Therefore, the findings indicate that radial symmetry types with alternating arrangements between successive rings are present throughout the body.

### 3.4 Radial symmetry type correlates with polyp diameter

How are the radial symmetry types in the tentacle arrangements selected? In plants, depending on the size of the meristem (undifferentiated stem cell tissue), there are differences in the number of leaf organs per whorl, demonstrating a positive correlation (Rutishauser, 1999). In Cnidaria, *Hydra* has multiple tentacles that are arranged in a ring with different numbers; which is shown to be positively correlated with polyp size, with polyps with a larger diameter having more tentacles (Parke, 1900). However, whether the polymorphism of the symmetric arrangement associated with organ number variations exists in a size-dependent manner remains unelucidated. In contrast to *Hydra*, *C. uchidai* polyps have multiple rings, and the tentacle number of each ring is inherited from the primary ring. Above we observed that the number of tentacles within the primary ring defines the symmetry type (Fig. 4). Therefore, we elucidated whether the symmetry type is dependent on the diameter of *C. uchidai* polyps in the primary ring area as this defines the radial symmetry type (Fig. 5A). Polyp diameter significantly increased linearly with the type of radial symmetry (tri, tetra, and penta) but not with the 3D Euclidean distances measured between adjacent tentacles within the primary ring (Fig. 5B). Taken together, our results indicate that size correlates not only with tentacle number within the ring but also with symmetry type, suggesting that polymorphisms in symmetries arise from the polyp diameter size variation.

**Figure 5.**
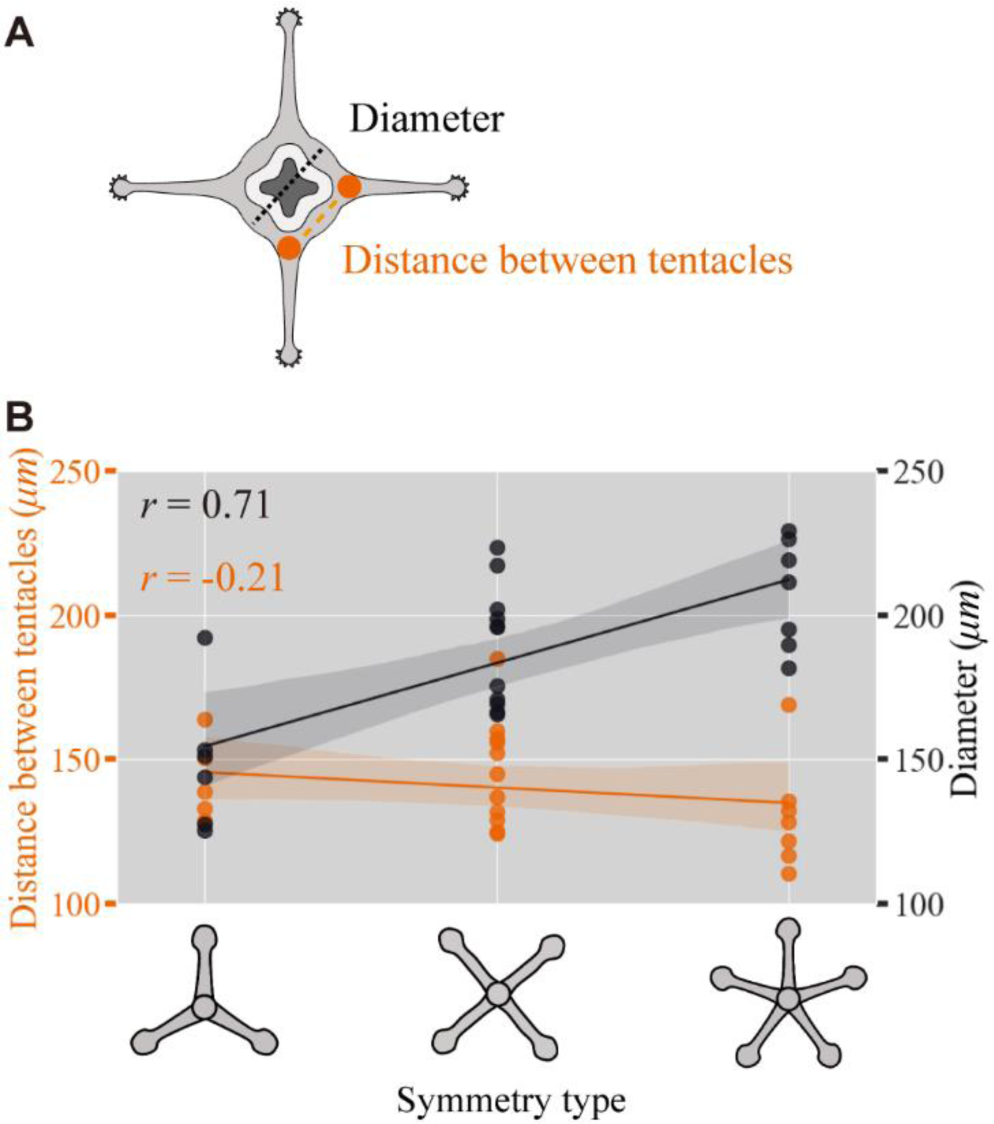
Polyp diameter-correlated radial symmetry type. **A** Schematic view of the polyp showing the diameter (black dotted lines) and distance (orange dashed lines) between the adjacent tentacles. **B** *C. uchidai* polyp diameter of individuals (black plot lines) and the average distance between adjacent tentacles measured in each individual (orange plot lines) as a function of the radial symmetry type corresponding to tri-, tetra-, and pentaradial symmetries with the Pearson’s correlation coefficient 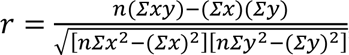, where *x*, *y,* and Σ denote the symmetry types, polyp diameter size or distance between tentacles, and the summation for the observed individuals (n = 24), respectively.

### 3.5 Mathematical model for tentacle arrangement

To predict the mechanism underlying periodic tentacle arrangements and size-correlated polymorphisms in radial symmetry, we built a mathematical model. Previous mathematical models for tentacle arrangements in *Hydra* (Meinhardt, 2012; Meinhardt, 1993) have suggested that molecular gradients decide the placement of the mouth, foot (aboral side), and a tentacle ring containing regularly spaced tentacles close to the mouth while maintaining a certain distance. These characters are consistent with our observations in *C. uchidai* polyps (Fig. 2C), whereas the *Hydra* models unexplored the following characteristics of *C. uchidai* polyps: radial symmetry throughout the body is evident in multiple periodic rings comprising regularly spaced tentacles and alternate arrangements established in successive rings (Figs. 2C-F, 3F, M, R and 4).

Therefore we developed our model based on a previous model on *Hydra* (Meinhardt, 2012). In the *Hydra* model, three diffusive substances were assumed to be involved in tentacle initiation: activator (A), inhibitor (B), and lateral inhibitor (C). The activator (A) and inhibitor (B) morphogens were assumed to be secreted from the mouth area, since tentacle initiation at the oral area while maintaining a certain distance to the mouth. Lateral inhibitor (C) inhibits tentacle initiation and is secreted from each tentacle to reproduce regular spacing in the tentacles of *Hydra*. Since the characteristics described in previous studies, namely, tentacle initiation at the oral area with a certain distance and regular spacing of the tentacles, are common in *C. uchidai*, we incorporated the previous settings that a cell in the polyp body region satisfying the following conditions became a tentacle: when the concentration of activator A was more than the threshold (*a* > *T_a_*; activator threshold in Fig. 6B) and the concentrations of inhibitors B and C were below the other thresholds (*b* < *T_i_*, inhibitor threshold; *c* < *T_ti_*, tentacle inhibitor threshold). However, the setting alone would produce multiple tentacles at approximately the same level, inconsistently with our observations (Figs. 2F, 3F). To this end, we additionally assume that the tentacle initiates at the global maximum of the activator.

**Figure 6.**
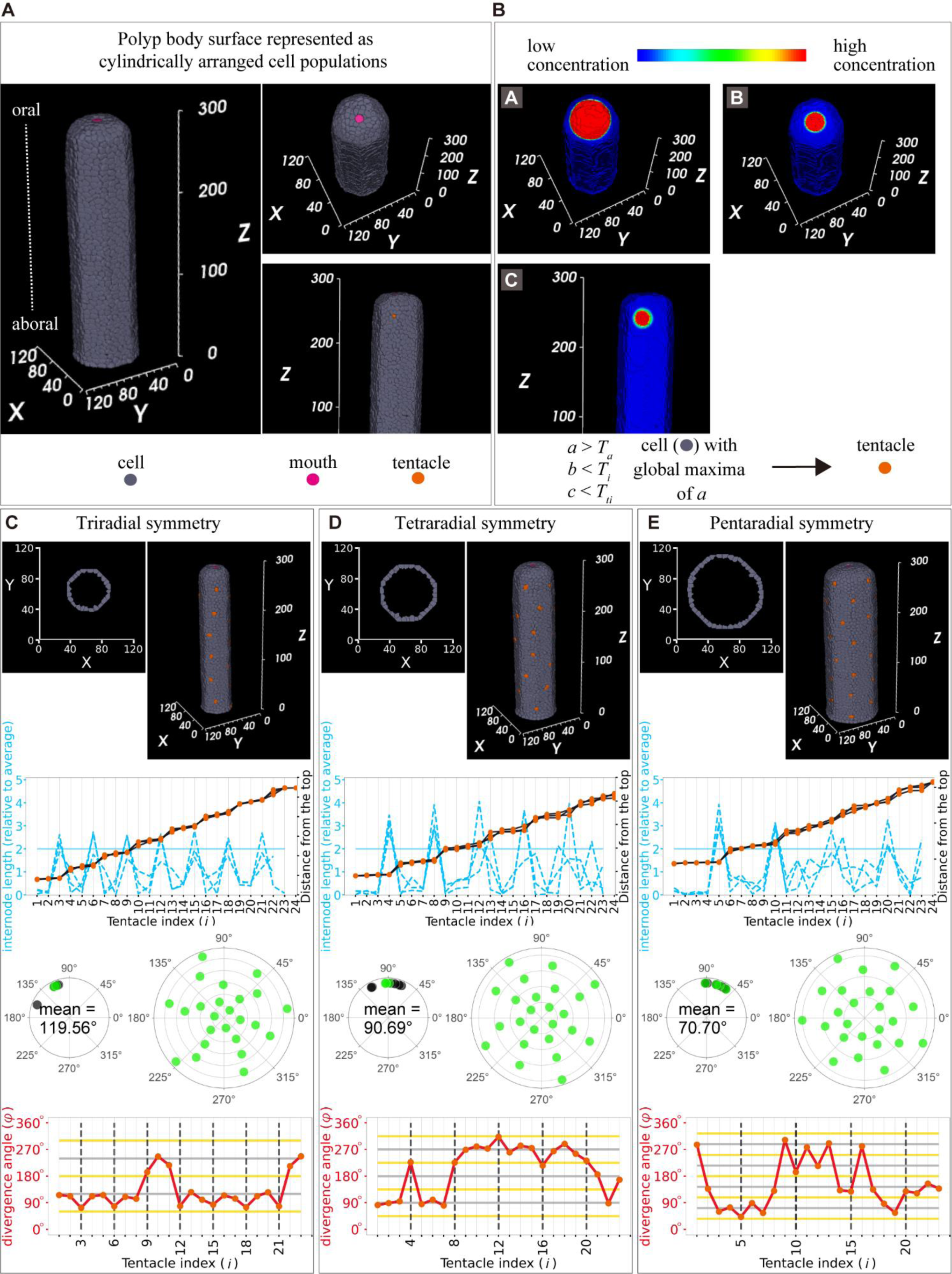
Mathematical model for tentacle arrangement. **A** Model settings: cylindrical polyp body, cells (grey), mouth (pink), and tentacle (orange). **B** Spatial patterns of the three morphogens: activator (*a*) **(B, upper left panel)** and two inhibitors (*b* and *c*) **(B, upper right panel and middle panel)**. All morphogens were assumed to diffuse in the single-cell layer (grey region) but not in the other layers. Simulated conditions of the concentrations of the activator and inhibitor **(lower panel)** in the cell (grey) till tentacle initiation (orange). *T_a_*, activator threshold; *T_i_*, inhibitor threshold; and *T_ti_*; tentacle inhibitor threshold. **C-E** Cells positioned cylindrically in the three-dimensional space and horizontal sections in two-dimensional space after simulations producing tentacles (at cells in orange in **top panel)**: tri-, tetra-, and pentaradial symmetries were reproduced at different diameters (54 in **C**, 72 in **D**, 90 in **E**). Internode length (blue dashed lines) of the successive tentacles and distance from the mouth (black plot lines) as a function of tentacle indices with tri- **(C)**, tetra- **(D)**, and pentaradial **(E)** symmetries in the second panels from the top ones. Angles measured based on the differences in the position angles in tri- **(C, third left panel from the top,** 119.56°±17.22°), tetra- **(D, third left panel from the top,** 90.69°±17.19°), and pentaradial **(G, third left panel from the top,** 70.70°±12.73°) symmetries in the primary ring in the representative sample and other samples indicated with green and black dots, respectively. Polar plots of the position angles (green) of the tentacles of each polyp with tri- **(C, third right panel from the top)**, tetra- **(D, third right panel from the top)**, and pentaradial **(E, third right panel from the top)** symmetries. Divergence angles (red lines) of the successive tentacles as a function of tentacle indices with tri- **(C, bottom panel)**, tetra- **(D, bottom panel),** and pentaradial **(E, bottom panel)** symmetries. The grey and yellow horizontal lines indicate multiples of period angles (120° in tri-, 90° in tetra-, and 72° in pentaradial symmetries) and their half, respectively. Black dashed vertical lines indicate the numbers for each period. (n = 3)

Because the *Hydra* model reproduces a single ring, we examined whether the inhibition and activation model accounts for the periodically formed multiple rings comprising tentacles alternately arranged in successive rings observed in *C. uchidai*. By performing model simulations on the single cell-layer surface of a cylindrical polyp body (Fig. 6A), the tentacles were initiated near the mouth while maintaining a certain distance (Fig. 6C-E, top panels) owing to the suprathreshold of activator *a* and subthreshold of inhibitor *b.* At an intermedium size of polyp diameter, four tentacles were sequentially initiated and regularly arranged in the primary ring, demonstrating a position angle difference between the nearest tentacles at ∼90° (90.69°±17.19°, n = 3) owing to inhibitor *c* (Fig. 6D). Tentacle initiations proceeded through the aboral area owing to activator secretion from the mouth (Supplemental Fig. S1). Since the activator is effective throughout the body and lateral inhibitor diffused from each tentacle, multiple rings formed with the periodicity in every four tentacles (*R*_4_ = 0.99, *p* = 5.6 × 10^−12^ in Fig. 6D bottom panel), while tentacles were alternately initiated in successive rings (Fig. 6C-E, top and lower panels). In particular, as the diameter of the polyp body increased, the number and angular arrangement of the tentacles within the primary ring selectively exhibited tri-, tetra-, and pentaradial symmetries (Fig. 6C-E). Therefore, the activation and inhibition of tentacle initiation reproduced the periodically formed multiple rings comprising regularly spaced tentacles, with different radial symmetries correlating with polyp size, as observed in *C. uchidai*. Taken together, our model suggests the regulation of size-oriented symmetry selection of organ arrangements in 3D space.

## 4. Discussion

### 4.1 Tentacle arrangement principles and radial symmetry selection

Our quantitative analysis revealed that ring arrangements were defined by internode lengths (Fig. 3F), resembling the whorled arrangement in plant phyllotaxis (Fig. 3D). The whorled arrangement in plants demonstrates constant divergence angles within the whorls (Fig. 3K). Consistently, regularly spaced tentacles within the ring were evident in the equal angles between the position angles (Fig. 3R). However, in tentacles, we could not find this consistency in divergence angles (Fig. 3M), while the divergence angles between successive rings revealed alternate arrangements in successive rings (Fig. 4E, F, G, lower panels). Therefore, tentacle arrangements in hydrozoan polyps are similar to those in whorled plant arrangements, and the phyllotaxis measurement system can be adapted for tentacle organ arrangements of hydrozoan polyps.

*Hydra* is the target of studies on tentacle arrangements forming a single ring because of the satisfactory amount of the activator in a limited area (Meinhardt, 2012). In contrast to *Hydra*, a considerable number of hydrozoan species form multiple rings throughout the polyp body, although whether similar mechanisms regulate tentacle arrangements remains unknown. A previous study has reported tentacle formation throughout the body column of *Hydra* after treatment with the drug alsterpaullone, which increased the amount of the activator (Meinhardt, 2012). Therefore, in the present model, we considered a sufficient amount of the activator for tentacle initiation throughout the polyp body, enabling multiple ring arrangements in the 3D space (Fig. 6C-E), consistent with those observed in *C. uchidai* (Fig. 2F). Future studies examining the molecular background of tentacle arrangements in hydrozoan polyps can clarify whether the amount of activator accounts for the transition from a single ring to multiple periodic rings, establishing the radial symmetry of organ arrangements in 3D space.

In some hydrozoan polyps, tentacle number variations have been reported (Beklemishev, 1969); however, quantitative analysis of the tentacle arrangement needed for symmetry identification was missing in most cases. In the present study, quantitative analysis revealed the principles of tentacle arrangement, including regularly spaced tentacles comprising periodic rings and alternations in successive rings (Fig. 3 and 4). Moreover, we found polymorphisms in tri-, tetra-, and pentaradial symmetries correlated with *C. uchidai* polyp size (Fig. 5B). Furthermore, our developed model reproduced these tentacle arrangements and size-dependent tentacle number decisions (Fig. 6C-E), suggesting the mechanisms underlying the size-oriented tentacle number variations, which selectively determine the type of radial symmetry. Similar to *C. uchidai*, other hydrozoan polyps exhibit simple cylindrical body structures, considered in the present model (Fig. 6A). Therefore, our established quantification methods (Fig. 3) and mathematical model for tentacle arrangements (Fig. 6) can be applied to other hydrozoan polyps, revealing if the size-dependent symmetry selection and periodic arrangement principles exist in hydrozoan ancestors and other radially symmetrical animals.

### 4.2 Size-dependent polymorphism

Our results reveal that tentacle arrangement, particularly the number of tentacles within a ring, depends on polyp body diameter, indicating a positive correlation between organ number and field size (Fig. 5B). In plants, phyllotactic pattern formation occurs at the periphery of the apical region, while the central zone is kept meristematic. Theoretical studies have reported that the phyllotactic pattern depends on the size of the central zone and that the number of organs within a whorl in a whorled arrangement can be increased by increasing the size of the central zone (Douady and Couder, 1996; Jönsson et al., 2006; Kitazawa and Fujimoto, 2015). This finding is supported by a molecular study that reported that an increase in the stem cell population in the floral meristem increases the number of floral organs and that a decrease in the population disrupts the ring-like arrangement of the Arabidopsis flower (Schoof et al., 2000). Pattern formation by diffusible or transportable molecules generally results in the Turing-like pattern with constant spacing if it satisfies simple requirements (Turing, 1952; Meinhardt, 1982; Kondo and Miura, 2010), and size dependence of the organ arrangement is a natural consequence of pattern formation.

What is the biological significance of tentacle arrangement and its polymorphisms? Traditionally, there are two different but consistent views on the biological significance of plant phyllotaxis. The first view emphasizes the adaptive significance of widespread phyllotactic patterns, for example, focusing on light capture efficiency (Strauss et al., 2019). Another view insists that the pattern is constrained by developmental processes because phyllotactic patterns are easily produced, even in non-biological processes (Douady and Couder, 1992). Similarly, our results can be discussed in terms of both adaptive significance and growth constraints. Feeding efficiency is one potential candidate for the selection pressure of tentacle arrangement. Further studies on the feeding behaviors of the polyps will improve our understanding of the relationship between symmetrical arrangements and feeding efficiency.

### 4.3 Model limitations

Live imaging revealed that tentacle initiations started from the oral area and proceeded through the aboral side simultaneously as the polyp grew longitudinally (Fig. 2F). In addition to longitudinal polyp growth, an increase in diameter was evident in late-stage polyps compared with early-stage polyps (data not shown). Quantitative analysis revealed that the rings were present near the mouth but were either absent or sometimes present as ring-like arrangements comprising different numbers of tentacles in areas distant from the mouth (Fig. 3F and 4A-C). These results suggest that some instabilities were incorporated into tentacle arrangements during growth, resulting in number variations, particularly in growing areas. The present model performed calculations on the cylindrical cell area representing the grown body and did not reproduce some samples, such as rings comprising numbers different from the primary ring. On the other hand, our simulations resulted in ring arrangements appearing multiple times in the triradial symmetry (Fig. 6C, second panel from the top) compared with polyp samples (Fig. 4A) that exhibited disturbances in the aboral area. Incorporating the increase in diameter during growth could solve these inconsistencies between the model and real samples, as phyllotaxis models have revealed that pattern transition can occur by increasing stem diameter (Douady and Couder, 1996b; Kwiatkowska and Florek-Marwitz, 2014; Zagórska-Marek and Szpak, 2008). Consistently, our size measurements and model revealed that polyp diameter correlates with tentacle numbers within the ring (Figs. 5 and 6C-E), suggesting that these number variations can arise because of fluctuations (growth speed and amount) during growth processes. Future studies incorporating growth processes that reproduce fluctuations in the model can clarify how number variations emerge in the successive rings.

## 5. Conclusion

We revealed the principles of tentacle arrangements that form periodic rings comprising multiple regularly spaced tentacles, establishing radial symmetry. Furthermore, we observed polymorphisms in the type of symmetry, including tri-, tetra-, and pentaradial symmetries, which are positively correlated with polyp diameter, with a larger diameter in pentaradial symmetry than in tetra- and triradial symmetries. Our established quantification methods and mathematical model for tentacle arrangements are applicable to other radially symmetrical animals, and will reveal the widespread association between the size-correlated symmetry and periodic arrangement principles.

## 6. Conflict of interest

The authors declare that the research was conducted in the absence of any commercial or financial relationships that could be construed as a potential conflict of interest.

## 7. Author contributions

Conceptualization, SS and KF; Data curation, SS; Formal analysis, SS and MK; Funding acquisition, SS and KF; Investigation, SS and NT; Methodology, SS and MK; Project administration, SS; Resources, SS; Software, SS; Supervision, SS; Validation, SS; Visualization, SS and MK; Writing— original draft, SS, MK, NT, and KF; Writing—review and editing, SS, MK, and KF. All authors contributed to the manuscript and approved the submitted version.

## 8. Funding

This work was supported by Grants-in-Aid for Scientific Research from the Ministry of Education, Culture, Sports, Science, and Technology of Japan for SS (22KJ3132) and KF (21K19297, 22H04719).

## 9. Acknowledgments

We would like to thank Dr. Shigeru Kuratani for giving their valuable advice regarding the discussions on animal symmetry.

## 11. Supplementary Material

Supplementary Figure S1.

Temporal evolution (represented by t) of the 3D arrangement of cells and tentacle initiations

Supplementary Table S1.

Parameters used in the mathematical model Supplementary Data S1

Raw data for tentacle position. The first line indicates the sample identifier, which follows the cylindrical coordinates of heights *h*_i_ and position angles θ*_i_* of each tentacle *i* ordered by height, and −1 indicates the end of the dataset for a sample.

Supplementary Data S2

CompuCell3D script for simulating the mathematical model.

## 12. Data availability statement

The raw data and script supporting the conclusions of this article are provided in the Supplementary Material.

## Notes

### Competing Interest Statement

The authors have declared no competing interest.

