## Supplementary Data S1 for "Size-Correlated Polymorphisms in Phyllotaxis-Like Periodic and Symmetric Tentacle Arrangements in Hydrozoan Polyps"

1.76.3

| | |
|---|---|
| 80.61078918 | 188.9732309 |
| 84.31500582 | 317.3160801 |
| 97.91553444 | 65.26965206 |
| 163.6067392 | 255.1812455 |
| 168.5355854 | 8.615707389 |
| 176.9251134 | 126.4253418 |
| 211.4033037 | 72.87717948 |
| 221.9191451 | 202.5381647 |
| 229.3574596 | 303.3188421 |

-1

1.76.4

| | |
|---|---|
| 76.4548004 | 171.5526255 |
| 87.595865 | 281.1229964 |
| 88.99150729 | 62.25307325 |
| 116.1408476 | 358.100999 |
| 139.4060379 | 116.4380853 |
| 146.9873923 | 224.0108936 |
| 167.7286031 | 293.0638995 |
| 168.2079579 | 168.0966253 |
| 169.9502007 | 46.12938059 |

-1

1.76.5

| | |
|---|---|
| 90.0874326 | 342.0742111 |
| 92.41076239 | 164.4296466 |
| 93.74703145 | 66.91259783 |
| 94.85438265 | 253.3714219 |
| 172.3985284 | 109.9763675 |
| 175.7285751 | 196.8378321 |
| 180.8132915 | 22.21536377 |
| 187.3549897 | 286.6555737 |

-1

1.76.6

| | |
|---|---|
| 72.74102126 | 250.0277996 |
| 93.02293565 | 158.916812 |
| 94.50641511 | 58.81414936 |
| 94.79892175 | 327.5908082 |
| 134.5482386 | 106.1875113 |
| 136.4690508 | 216.0597626 |
| 147.2439123 | 8.994052951 |
| 152.1329744 | 289.8813433 |

-1

1.76.7

| | |
|---|---|
| 85.27541332 | 270.1416395 |
| 88.59282145 | 145.5470412 |
| 100.9668016 | 36.47407618 |
| 148.9158978 | 221.9000842 |
| 152.1406391 | 330.6530915 |
| 157.661427 | 94.78078517 |
| 190.159589 | 268.5343594 |
| 195.0800889 | 147.4141606 |
| 207.4534155 | 36.23467488 |

-1

1.76.8

| | |
|---|---|
| 90.08586986 | 216.2743029 |
| 92.16456341 | 83.07431819 |
| 92.55732117 | 320.983205 |
| 140.9299553 | 155.4556432 |
| 154.0252842 | 20.3560236 |
| 157.5960663 | 271.1425417 |
| 180.2396916 | 102.4588878 |
| 186.2338668 | 201.8552414 |
| 198.2409562 | 322.1729457 |

-1

1.77.1

| | |
|---|---|
| 62.04075893 | 348.6890264 |
| 73.63225475 | 136.4807736 |
| 78.12102674 | 208.1481364 |
| 97.60013591 | 305.6181739 |
| 108.0187482 | 48.84018034 |
| 110.6927601 | 171.304392 |
| 115.9364267 | 352.4060362 |
| 131.3438757 | 228.8407161 |
| 137.0005351 | 136.562427 |

-1

1.77.2

| | |
|---|---|
| 80.68137902 | 130.9758546 |
| 92.5376179 | 207.7603504 |
| 94.19182335 | 37.31311059 |
| 96.42492512 | 305.5928985 |
| 126.8538001 | 165.8810467 |
| 135.064245 | 258.6178286 |
| 136.825193 | 349.1822979 |
| 137.2930196 | 79.56677873 |

−1
1.77.3

| | |
|---|---|
| 117.7964776 | 320.0374507 |
| 125.319472 | 69.3500197 |
| 131.4722485 | 167.2459765 |
| 145.8857629 | 242.6892769 |
| 210.8308415 | 13.44919269 |
| 223.1911087 | 119.3288854 |
| 225.6428295 | 293.1518575 |
| 230.4019252 | 196.2809343 |
| 259.7485409 | 250.3403764 |
| 262.6422472 | 337.8125783 |
| 267.0940242 | 54.07313938 |
| 267.2105492 | 142.6284632 |

−1
1.77.4

| | |
|---|---|
| 97.35363822 | 348.8789668 |
| 108.7879192 | 160.5091725 |
| 109.6882999 | 57.96424288 |
| 115.7949929 | 239.7505304 |
| 154.5627792 | 293.3788573 |
| 164.2192695 | 28.29253177 |
| 183.047336 | 103.4339716 |
| 186.3976226 | 340.82992 |
| 188.4078347 | 193.5046548 |
| 198.9382407 | 56.03910189 |
| 208.8366155 | 237.298897 |
| 224.9625336 | 274.2607994 |
| 226.1467819 | 23.69539673 |
| 236.9205008 | 158.5370973 |
| 252.8849853 | 84.33768115 |
| 261.0407007 | 219.4656813 |

−1
1.77.5

| | |
|---|---|
| 80.42256734 | 192.2940529 |
| 85.45597254 | 327.8654415 |
| 89.29258346 | 80.04368825 |
| 119.9250967 | 255.3868707 |
| 127.1180769 | 23.79930121 |
| 153.3619104 | 141.4705807 |
| 154.4088573 | 316.0187087 |
| 164.3532976 | 204.948591 |

| | |
|---|---|
| 169.9071055 | 71.87824464 |
| -1 | |
| 1.77.7 | |
| 114.4588077 | 321.4925637 |
| 122.9576224 | 67.34027877 |
| 131.2967772 | 168.0893735 |
| 139.5492784 | 241.3020635 |
| 211.1162397 | 10.05479385 |
| 221.3939337 | 119.6857545 |
| 226.8001814 | 296.0438231 |
| 230.8643279 | 195.1660122 |
| 259.6209115 | 340.0434826 |
| 260.7196783 | 248.0819842 |
| 266.9476679 | 58.28000346 |
| 267.06383 | 145.8184662 |
| -1 | |
| 1.77.8 | |
| 112.2142857 | 56.13892612 |
| 120.0778951 | 180.5745514 |
| 134.1133497 | 302.3764882 |
| 172.4019349 | 135.1560424 |
| 177.4541582 | 358.205209 |
| 201.0517005 | 229.6643581 |
| 214.6094037 | 44.30016025 |
| 234.4890656 | 177.5675626 |
| 237.5291068 | 312.4650524 |
| 263.9396367 | 107.7576798 |
| 268.5312409 | 2.247639781 |
| 275.8324956 | 233.233045 |
| 287.278579 | 322.2362759 |
| 287.9360804 | 174.1783889 |
| 294.7251562 | 39.43724546 |
| -1 | |
| 1.77.9 | |
| 101.7565414 | 174.2270847 |
| 105.736871 | 259.5181666 |
| 107.0034255 | 349.1742647 |
| 111.0653891 | 75.16473369 |
| 160.463627 | 218.7586604 |
| 162.7443002 | 96.19790386 |
| 171.2346833 | 31.61243694 |
| 173.6338 | 306.7711704 |

| | |
|---|---|
| 187.4366978 | 159.7244477 |
| 188.1686076 | 262.456915 |
| 211.9367829 | 354.0498087 |
| 212.2962642 | 77.41700107 |
| -1 | |
| 1.77.10 | |
| 105.1001508 | 359.8779785 |
| 117.0783368 | 273.5180652 |
| 121.3076509 | 86.92845033 |
| 122.5190816 | 173.2616006 |
| 194.575583 | 42.40077476 |
| 196.0903885 | 311.6015509 |
| 202.8215293 | 140.1507521 |
| 209.9455612 | 227.7001602 |
| 228.5667364 | 96.03647074 |
| 231.3538603 | 359.5207414 |
| 235.9400264 | 189.9813614 |
| 238.8783068 | 263.2254602 |
| -1 | |
| 1.77.11 | |
| 117.4795401 | 132.5194789 |
| 118.4870603 | 271.7338344 |
| 135.648366 | 28.15684671 |
| 167.2455374 | 197.1814728 |
| 173.8472504 | 309.8773181 |
| 186.9627786 | 96.08590792 |
| 251.589288 | 14.05310376 |
| 253.5656082 | 130.5920082 |
| 256.9842186 | 272.4498442 |
| 286.5831757 | 93.50998193 |
| 294.246635 | 183.7158735 |
| 299.8997907 | 306.6670507 |
| 309.9005862 | 128.9227369 |
| 310.0113373 | 12.55881307 |
| 315.8241207 | 271.3920889 |
| -1 | |
| 1.79.1 | |
| 101.5291724 | 34.46394714 |
| 110.3833323 | 267.5240733 |
| 112.9249637 | 160.7795879 |
| 151.285041 | 85.39005864 |
| 153.7729956 | 219.0354182 |

| | |
|---|---|
| 162.9587816 | 332.2890778 |
| 205.3825047 | 39.54021129 |
| 212.7130554 | 262.3488961 |
| 219.2606647 | 157.2676065 |
| 244.9308131 | 318.4773945 |
| 249.3512357 | 223.9341584 |
| 251.8426938 | 84.63563488 |
| 265.4425829 | 33.01838675 |
| 267.6058503 | 262.1856529 |
| 272.8101039 | 169.9504616 |
| -1 | |
| 1.79.2 | |
| 104.6710995 | 327.8878239 |
| 112.9771472 | 213.2560368 |
| 116.2827567 | 84.66976385 |
| 148.3375949 | 276.3199802 |
| 155.3709066 | 35.49327222 |
| 163.7081925 | 147.3206896 |
| 194.5935443 | 335.1939335 |
| 228.9126178 | 209.2377867 |
| 236.6107045 | 97.70587939 |
| 261.840513 | 37.43748839 |
| 263.342904 | 270.9477101 |
| 283.4668834 | 146.1943369 |
| 289.7317405 | 332.8246132 |
| 293.4947027 | 225.7468709 |
| 303.1086537 | 86.70958917 |
| -1 | |
| 1.79.3 | |
| 100.1778258 | 274.5284043 |
| 104.1153113 | 13.8544716 |
| 107.4415869 | 113.2393666 |
| 114.3203939 | 181.5045392 |
| 173.1086422 | 341.0709608 |
| 178.8088509 | 214.0249621 |
| 190.8630055 | 53.45141592 |
| 199.6551361 | 150.4308441 |
| 207.1457973 | 285.9246772 |
| 220.9868171 | 8.64245163 |
| 234.9838645 | 189.4261818 |
| 238.1040374 | 96.98572725 |
| 239.2510346 | 258.0503748 |

| | |
|---|---|
| 254.3314766 | 152.3416547 |
| 257.0278014 | 337.667262 |
| 257.5876851 | 30.30438374 |
| -1 | |
| 1.79.4 | |
| 109.2181321 | 171.1915895 |
| 110.4804269 | 284.6704888 |
| 133.4020902 | 51.44358246 |
| 188.838175 | 223.4013176 |
| 207.5318825 | 103.7119426 |
| 212.860698 | 341.3481444 |
| 227.375459 | 165.9526828 |
| 244.7385352 | 276.1316457 |
| 255.1761653 | 42.87632966 |
| 264.5563577 | 219.1359005 |
| 267.5433588 | 118.3122664 |
| 286.2576027 | 340.3333629 |
| -1 | |
| 1.79.5 | |
| 116.8598407 | 336.217488 |
| 120.8036031 | 218.3613509 |
| 124.2724607 | 110.0111756 |
| 151.8009183 | 38.67500943 |
| 182.9436896 | 164.9950796 |
| 183.2232317 | 281.031798 |
| 214.6500256 | 349.4773343 |
| 242.0186614 | 112.9074131 |
| 248.4892002 | 216.099528 |
| 284.5428958 | 293.5982596 |
| 295.897442 | 41.9185144 |
| 318.8998893 | 160.9004924 |
| 334.5460832 | 344.728784 |
| 352.7464779 | 220.2875418 |
| 378.7487083 | 106.7880446 |
| 380.1555169 | 288.7511041 |
| 385.8718709 | 18.20760002 |
| 420.8123854 | 175.9952466 |
| -1 | |
| 1.86 | |
| 180.1347611 | 236.1470545 |
| 183.9075765 | 150.3400323 |
| 189.879689 | 49.79039076 |

| | |
|---|---|
| 196.5303329 | 318.003149 |
| 342.888942 | 97.05603692 |
| 355.3765941 | 358.3515473 |
| 360.7556523 | 269.0770033 |
| 384.5375232 | 186.7189649 |
| 538.3083173 | 50.1045591 |
| 560.4345691 | 306.8437473 |
| 563.6853117 | 218.6822397 |
| 616.7685026 | 136.0958923 |
| 742.1870502 | 0.1443189702 |
| 760.6984149 | 82.3890632 |
| 760.925891 | 270.5066176 |
| 798.113283 | 172.2224699 |
| 908.8174909 | 320.5067914 |
| 921.7455802 | 227.0517723 |
| 936.7298316 | 118.9430337 |
| 969.318988 | 37.92690653 |
| -1 | |
| 1.80.2n | |
| 136.144029 | 1.536799111 |
| 136.9258458 | 234.9920417 |
| 157.0414552 | 133.0109789 |
| 173.5610917 | 292.390056 |
| 188.8290822 | 71.64209405 |
| 217.2598642 | 177.9648675 |
| 247.0672769 | 14.04780032 |
| 282.5285998 | 241.2183888 |
| 298.6663139 | 113.9977363 |
| 328.8271374 | 309.7996496 |
| 369.0372779 | 176.0302746 |
| 374.5935903 | 56.54512811 |
| 405.7325006 | 5.302048558 |
| 420.1199952 | 252.5192644 |
| 432.5148929 | 112.5272078 |
| 468.4357165 | 58.72383596 |
| 468.46641 | 309.7190234 |
| 469.3547483 | 196.7559255 |
| 514.3750433 | 242.1908836 |
| 515.0001231 | 354.5942891 |
| 524.9826549 | 143.779732 |
| -1 | |
| 1.2 | |

| | |
|---|---|
| 214.6661004 | 128.1876174 |
| 237.0393656 | 210.8154355 |
| 261.2109509 | 324.8619829 |
| 275.0036071 | 48.33710271 |
| 396.2021145 | 269.4810646 |
| 473.8998641 | 179.3186956 |
| 501.309964 | 76.6090603 |
| 531.6936324 | 9.042038749 |
| 613.0358782 | 204.4555365 |
| 668.3834081 | 305.6815836 |
| 858.3445981 | 252.7877069 |
| 864.9737755 | 101.4646766 |
| 888.0842321 | 39.14744817 |
| 982.117401 | 164.7543927 |
| 989.8895036 | 337.464601 |
| 1144.537597 | 65.05066178 |
| 1149.376719 | 286.4844987 |
| 1177.12407 | 212.3776295 |
| 1231.61212 | 1.052992851 |
| 1245.063388 | 127.894765 |
| 1283.132518 | 54.51752359 |
| 1312.373454 | 337.1154103 |
| 1318.318527 | 254.2304405 |
| 1320.957982 | 177.7371786 |
| -1 | |
| 1.3 | |
| 192.3979504 | 355.2448974 |
| 194.1460167 | 66.27773978 |
| 206.2888539 | 278.848015 |
| 210.7628227 | 148.2232587 |
| 222.1154283 | 199.0902186 |
| 376.8054613 | 241.2678153 |
| 395.9017413 | 105.8575208 |
| 420.9374909 | 314.8094555 |
| 424.0222417 | 177.189513 |
| 429.5693556 | 26.62503886 |
| 715.1444917 | 56.02716238 |
| 720.0277912 | 271.6571882 |
| 749.5624645 | 349.3785773 |
| 765.6144218 | 131.6296968 |
| 804.9916828 | 209.9329594 |
| 1006.800524 | 303.1860938 |

| | |
|---|---|
| 1067.904756 | 174.7299355 |
| 1087.686235 | 18.17781774 |
| 1132.851845 | 93.99462545 |
| 1179.391297 | 240.2358619 |
| 1276.328013 | 330.5033637 |
| 1329.598079 | 144.7074893 |
| 1336.074684 | 48.81414364 |
| 1368.700502 | 231.2668509 |
| -1 | |
| 1.13 | |
| 141.9276615 | 348.001507 |
| 196.5915272 | 66.91509193 |
| 200.3654537 | 154.0246256 |
| 207.5878658 | 244.7046844 |
| 342.8330491 | 211.4198404 |
| 379.374922 | 305.932919 |
| 418.3756934 | 31.74949633 |
| 425.5249743 | 129.0557325 |
| 590.5378612 | 255.4812932 |
| 633.5486178 | 193.9822316 |
| 685.3783654 | 10.17618837 |
| 729.1445484 | 124.0372882 |
| 877.2622759 | 353.0352002 |
| 880.8432466 | 267.8756838 |
| 928.9487011 | 192.084395 |
| 961.2436646 | 101.2241524 |
| 1067.296878 | 338.8417782 |
| 1101.631243 | 248.510452 |
| 1122.41119 | 55.36206534 |
| 1130.909662 | 169.3670946 |
| 1201.13227 | 220.6360444 |
| 1202.302226 | 314.3711104 |
| 1206.838446 | 116.5838396 |
| 1242.259384 | 27.75439512 |
| -1 | |
| 1.14 | |
| 119.1046122 | 142.0802015 |
| 122.7694145 | 60.96604033 |
| 122.7729928 | 296.7813313 |
| 138.4747718 | 211.8692941 |
| 155.8291682 | 358.7723027 |
| 239.7857422 | 97.96234592 |

| | |
|---|---|
| 250.3603072 | 27.2302322 |
| 259.615581 | 178.9807835 |
| 288.8872958 | 270.9040955 |
| 330.2479964 | 342.8214597 |
| 369.5446794 | 141.6201121 |
| 384.3486913 | 219.8332389 |
| 395.6074271 | 73.31845011 |
| 430.823596 | 15.10105057 |
| 434.2209067 | 180.0715137 |
| 437.3384475 | 299.8589099 |
| −1 | |
| 1.15.2 | |
| 195.9796222 | 71.54200015 |
| 210.1884553 | 269.3896501 |
| 211.2123881 | 177.1300403 |
| 216.4953211 | 347.1326183 |
| 345.712377 | 221.7675012 |
| 354.7484696 | 124.8417179 |
| 381.0861982 | 37.00800991 |
| 409.7610892 | 306.6963206 |
| 552.2132504 | 173.6725313 |
| 581.7252267 | 83.1473516 |
| 589.3063097 | 352.4063555 |
| 628.7549188 | 260.4235452 |
| 822.118377 | 41.82600364 |
| 865.0683087 | 132.4812497 |
| 867.7230888 | 220.8308536 |
| 930.3890493 | 306.0346012 |
| 1037.797649 | 74.12004789 |
| 1047.519162 | 178.3855845 |
| 1050.609827 | 2.849176318 |
| 1058.568616 | 256.906463 |
| 1106.117356 | 46.71056256 |
| 1108.412171 | 126.2690741 |
| 1120.99062 | 218.9938061 |
| 1128.969381 | 304.6473356 |
| −1 | |
| 1.6 | |
| 190.9031189 | 166.1226112 |
| 195.0842482 | 236.5424108 |
| 201.5670505 | 83.95426886 |
| 208.6861386 | 324.7953108 |

| | |
|---|---|
| 210.7180051 | 23.68861644 |
| 338.9563147 | 265.1861767 |
| 351.8968457 | 138.3341501 |
| 388.0774386 | 203.3575825 |
| 410.7478648 | 351.3130725 |
| 436.3090131 | 59.2493229 |
| 620.6331329 | 303.2375226 |
| 665.1715348 | 227.395528 |
| 665.6117391 | 97.77793345 |
| 690.0788336 | 13.51029609 |
| 737.6777882 | 160.4038577 |
| 907.5010162 | 271.4780825 |
| 915.6622077 | 51.27242283 |
| 974.9735612 | 340.2275066 |
| 1043.625668 | 198.1318154 |
| 1079.267487 | 125.3986059 |
| 1186.512487 | 26.04691903 |
| 1218.379683 | 242.6510846 |
| 1223.230275 | 304.012371 |
| 1224.507206 | 72.68461075 |
| 1260.576003 | 157.0322271 |
| 1320.310998 | 217.6173025 |
| -1 | |
| 1.16 | |
| 181.2497992 | 197.9335659 |
| 189.2624444 | 90.86092705 |
| 192.5411722 | 351.5308519 |
| 215.0481007 | 270.8607814 |
| 308.9456554 | 144.727882 |
| 377.5504607 | 34.06143449 |
| 413.6490606 | 240.2799417 |
| 477.9166435 | 313.9877051 |
| 491.0118203 | 187.3468593 |
| 495.5217361 | 92.51073308 |
| 654.6184851 | 275.3058423 |
| 700.0432335 | 133.702835 |
| 704.5932149 | 50.27069974 |
| 753.5653194 | 349.2940541 |
| 792.3646057 | 221.4809379 |
| 883.9988768 | 85.62979508 |
| 903.5169304 | 175.0972595 |
| 903.8025339 | 307.9689002 |

| | |
|---|---|
| 977.479806 | 21.91271974 |
| 988.2316096 | 116.4968916 |
| 990.0639683 | 256.3024796 |
| 1003.992542 | 205.5514511 |
| 1024.352016 | 64.08906874 |
| 1062.759825 | 327.3877726 |
| −1 | |
| 1.17 | |
| 123.0360297 | 8.685430841 |
| 123.1165841 | 129.0054917 |
| 129.6615102 | 280.0548658 |
| 132.1441628 | 204.0405545 |
| 190.0322362 | 75.24158325 |
| 211.5708718 | 312.0769145 |
| 235.59998 | 182.9588258 |
| 238.6471702 | 239.1619583 |
| 252.8805993 | 23.52103265 |
| 264.4664521 | 127.0376831 |
| 295.9582116 | 76.52226 |
| 303.4189207 | 276.1169726 |
| 323.2784902 | 320.3911572 |
| 327.2113152 | 207.2740965 |
| 328.7359654 | 141.8400287 |
| 334.2797862 | 26.52907779 |
| 344.5616072 | 90.46870019 |
| 356.0511959 | 266.1584038 |
| −1 | |
| 1.18 | |
| 173.0529086 | 351.5246268 |
| 188.0819611 | 84.3348623 |
| 198.3469166 | 269.2487168 |
| 200.4886836 | 196.5829265 |
| 313.1192381 | 149.8932002 |
| 372.1709441 | 29.94447496 |
| 414.9277629 | 231.9165939 |
| 465.8942245 | 314.1242582 |
| 489.8185159 | 97.21222292 |
| 503.2974828 | 182.477478 |
| 647.4788949 | 272.8041354 |
| 699.4918835 | 140.9999572 |
| 702.5116991 | 50.78597705 |
| 743.3527908 | 354.5191812 |

| | |
|---|---|
| 785.2727199 | 219.7670967 |
| 898.6230929 | 85.79137979 |
| 903.8188384 | 308.6539398 |
| 906.0378567 | 171.7920722 |
| 974.8433795 | 22.97387593 |
| 986.704953 | 123.7937789 |
| 989.4369451 | 252.1400364 |
| 1008.616935 | 202.0069901 |
| 1032.683375 | 55.19979499 |
| 1049.054333 | 327.7617932 |
| -1 | |
| 1.19 | |
| 186.5210607 | 165.1055482 |
| 193.2492595 | 5.357611552 |
| 193.2755453 | 281.5066343 |
| 213.4810019 | 90.47127365 |
| 309.2545577 | 199.9097837 |
| 377.0975551 | 336.9536064 |
| 407.0067946 | 134.1893061 |
| 456.5914282 | 23.80393753 |
| 462.5306784 | 173.8855697 |
| 481.5946742 | 258.1447103 |
| 592.3650475 | 72.44973236 |
| 625.2379323 | 210.8964727 |
| 646.428917 | 316.5658338 |
| 684.2360577 | 1.160690763 |
| 731.9643723 | 143.5210032 |
| 821.4437045 | 273.0375971 |
| 841.575087 | 181.5308994 |
| 846.7095943 | 34.80426896 |
| 920.3759242 | 344.4145848 |
| 923.2770487 | 106.3849819 |
| 927.4785554 | 220.5424207 |
| 967.3051524 | 158.3579344 |
| 975.7794492 | 25.18583088 |
| 980.4849224 | 315.8357158 |
| -1 | |
| 1.22 | |
| 165.8983492 | 288.5623368 |
| 175.0950028 | 30.41244877 |
| 178.5175652 | 165.1842783 |
| 246.7536424 | 233.755594 |

| | |
|---|---|
| 252.6312087 | 328.2565015 |
| 258.0865022 | 97.72831437 |
| 320.2363617 | 163.4183462 |
| 325.463582 | 30.29187487 |
| 369.3251735 | 275.4763751 |
| 457.0325295 | 208.9778055 |
| 461.0089978 | 99.20287256 |
| 489.1643861 | 335.3094507 |
| 605.408709 | 162.0893209 |
| 620.5041709 | 49.89847704 |
| 634.6866031 | 277.5899958 |
| 773.1426673 | 352.7531225 |
| 777.3845488 | 114.1646876 |
| 780.2266143 | 239.5547022 |
| 898.9372387 | 297.3541884 |
| 917.7645316 | 176.848642 |
| 929.3142301 | 55.63644776 |
| 972.5824702 | 0.6631749146 |
| 973.5113634 | 242.2118532 |
| 1012.95361 | 122.2027597 |
| -1 | |
| 1.23 | |
| 213.5712332 | 207.6240454 |
| 224.1789662 | 150.974872 |
| 225.4735438 | 71.36986436 |
| 232.4032365 | 358.0767381 |
| 253.1546577 | 286.6668728 |
| 396.8323453 | 356.2218689 |
| 437.6047464 | 181.7058836 |
| 452.0923612 | 112.0636452 |
| 518.8614235 | 43.83192596 |
| 524.6234251 | 241.8742207 |
| 576.8424263 | 311.8532173 |
| 700.3820325 | 142.8886736 |
| 745.2469716 | 12.5451453 |
| 778.4704482 | 201.0532513 |
| 809.5963896 | 274.6245075 |
| 829.2536261 | 69.53437879 |
| 917.0236774 | 339.8182784 |
| 993.4825804 | 161.8462767 |
| 1015.520423 | 26.36040963 |
| 1057.393517 | 212.2922349 |

| | |
|---|---|
| 1090.134087 | 103.9993092 |
| 1134.270183 | 302.6655814 |
| 1216.388307 | 348.4706462 |
| 1231.743024 | 174.6088825 |
| 1265.336772 | 225.4821363 |
| 1281.460755 | 40.45794622 |
| 1302.664306 | 125.7874007 |
| 1346.380777 | 306.6478662 |
| 1362.742498 | 190.0336244 |
| 1395.001815 | 8.871949154 |
| -1 | |
| 1.24 | |
| 134.9910152 | 203.5603861 |
| 158.5693102 | 310.3501543 |
| 163.1505884 | 116.0331541 |
| 175.6637156 | 26.52152642 |
| 278.5292283 | 254.5153315 |
| 292.3439286 | 167.4393698 |
| 333.5568999 | 59.67807612 |
| 343.2953872 | 345.0633845 |
| 428.8896858 | 210.4550035 |
| 457.3392382 | 131.7073592 |
| 476.9876814 | 23.15315944 |
| 499.7895556 | 292.6820096 |
| 569.464048 | 178.1426123 |
| 579.2876521 | 89.96027697 |
| 604.4653241 | 340.5256946 |
| 619.3762692 | 249.3079733 |
| 626.5924542 | 33.27695468 |
| 648.4563633 | 150.3622232 |
| 676.6267036 | 302.0910172 |
| 678.0647911 | 202.0330817 |
| -1 | |
| 1.25 | |
| 171.670018 | 309.5371047 |
| 207.0183075 | 172.3987171 |
| 218.177263 | 41.70684638 |
| 287.9994266 | 225.5189426 |
| 305.9953758 | 108.702703 |
| 333.8177522 | 6.036402218 |
| 455.1519257 | 172.5155851 |
| 469.0305411 | 294.3929003 |

| | |
|---|---|
| 524.6513261 | 47.5972423 |
| 613.9831247 | 113.7838242 |
| 623.4539978 | 225.5877151 |
| 641.5072842 | 359.1903787 |
| 762.2334966 | 179.8087354 |
| 799.9883384 | 39.08099032 |
| 804.1136091 | 299.6146811 |
| 985.3190518 | 88.02253857 |
| 1002.063553 | 341.9505968 |
| 1022.146073 | 145.9954048 |
| 1081.272497 | 245.318827 |
| 1258.371775 | 18.77589405 |
| 1300.768502 | 210.1051359 |
| 1375.45974 | 309.0043695 |
| 1460.861279 | 63.2848441 |
| 1523.528354 | 186.1349807 |
| 1577.739331 | 359.0368693 |
| 1620.185088 | 252.9700194 |
| 1620.671843 | 133.3760985 |
| 1646.784888 | 316.2745497 |
| 1677.884928 | 57.21488373 |
| 1689.315581 | 205.0673439 |
| -1 | |
| 1.27 | |
| 181.7235066 | 116.1432967 |
| 194.3405727 | 208.4904043 |
| 200.4242884 | 38.60145236 |
| 204.9473788 | 305.8812389 |
| 291.8314805 | 256.5202953 |
| 385.9751711 | 162.0341005 |
| 389.8472662 | 1.439967719 |
| 397.3132575 | 69.15063315 |
| 489.0073294 | 218.2185824 |
| 490.3020314 | 286.8325222 |
| 622.39375 | 114.0858728 |
| 636.6426803 | 27.96287593 |
| 701.3618813 | 249.6051279 |
| 725.7831526 | 329.1657183 |
| 734.5367229 | 177.0110442 |
| 735.1163576 | 59.47002641 |
| 873.0561726 | 130.4242179 |
| 875.530366 | 220.5482878 |

| | |
|---|---|
| 881.1378771 | 76.74868769 |
| 887.8801122 | 282.1974888 |
| 905.430903 | 26.20392469 |
| 954.3167356 | 178.2737168 |
| 982.0437961 | 328.6212511 |
| 1003.247203 | 248.492888 |
| 1016.253409 | 55.50454844 |
| 1022.65532 | 112.7515628 |
| 1023.285452 | 297.2438967 |
| 1024.804006 | 9.549019276 |
| -1 | |
| 1.28 | |
| 198.0623069 | 99.7875786 |
| 206.1391123 | 267.5121381 |
| 212.1299406 | 183.4308206 |
| 219.0353815 | 4.613437628 |
| 361.786347 | 318.8006172 |
| 376.7696032 | 50.80628861 |
| 413.9058927 | 222.9283288 |
| 423.7897753 | 143.1196435 |
| 532.373139 | 8.129196643 |
| 572.7912523 | 274.0772537 |
| 616.27156 | 174.1572455 |
| 634.2379929 | 87.42590018 |
| 729.2264037 | 323.2158284 |
| 787.1343575 | 231.9914716 |
| 804.583832 | 25.27876176 |
| 862.7408952 | 132.274859 |
| 977.6892704 | 268.7377145 |
| 991.8433832 | 184.247952 |
| 1017.146054 | 83.3678374 |
| 1031.27143 | 346.3037701 |
| 1055.74221 | 231.5618816 |
| 1121.540752 | 128.0152558 |
| 1158.109703 | 298.9179707 |
| 1176.216417 | 41.48105862 |
| 1189.554681 | 192.840812 |
| 1203.258017 | 257.2262501 |
| 1214.499453 | 355.7904033 |
| 1232.861367 | 77.63754029 |
| -1 | |
| 1.30.1 | |

| | |
|---|---|
| 194.0991736 | 204.2720853 |
| 208.0227046 | 97.70989865 |
| 215.4471114 | 292.1089193 |
| 216.2189192 | 13.71523259 |
| 315.8747505 | 160.452071 |
| 471.2847452 | 244.2987413 |
| 475.5488268 | 342.0272749 |
| 485.9707784 | 64.27229999 |
| 597.7828584 | 123.5721364 |
| 628.3101164 | 293.6861879 |
| 649.4883282 | 190.2409929 |
| 721.226391 | 20.45412597 |
| 806.8345966 | 85.25924558 |
| 845.2363442 | 256.6557036 |
| 853.5125871 | 329.4307831 |
| 881.2158053 | 149.2329248 |
| 933.8553771 | 32.16422115 |
| 945.9192611 | 286.0258674 |
| 956.993334 | 204.2930383 |
| 990.3839602 | 152.5990178 |
| 993.7424204 | 93.9103763 |
| 993.8744326 | 253.8518107 |
| 999.5117548 | 342.7439909 |
| -1 | |
| 1.30.2 | |
| 202.027708 | 40.91022892 |
| 217.8413976 | 134.3012617 |
| 223.3953716 | 315.0874355 |
| 241.9398343 | 223.9860585 |
| 430.7482159 | 80.74846106 |
| 450.3178544 | 353.1472934 |
| 497.7208878 | 254.6635638 |
| 532.1218878 | 179.0816241 |
| 728.2656092 | 119.4296029 |
| 772.7621388 | 303.7983863 |
| 808.6866297 | 23.12091696 |
| 911.5818646 | 216.1868409 |
| 1040.853415 | 74.76599198 |
| 1108.813525 | 346.5800911 |
| 1120.0526 | 161.0468874 |
| 1224.949343 | 272.740788 |
| 1358.207378 | 47.56430793 |

| | |
|---|---|
| 1398.003682 | 219.9410733 |
| 1439.490738 | 118.0889843 |
| 1520.202891 | 338.9583351 |
| 1579.293996 | 174.9435886 |
| 1613.50774 | 79.45725041 |
| 1659.156327 | 122.4222634 |
| 1659.450079 | 13.80844457 |
| 1662.513469 | 281.6275377 |
| 1671.82906 | 212.8800936 |
| 1716.922342 | 44.44910773 |
| 1751.978592 | 325.4314748 |
| -1 | |
| 1.31 | |
| 203.5891987 | 92.92597697 |
| 209.6346576 | 274.9929458 |
| 218.2308854 | 356.7336674 |
| 234.943686 | 182.1138639 |
| 360.450314 | 39.09061427 |
| 404.3354462 | 225.2077991 |
| 442.5769038 | 146.1252449 |
| 465.3642528 | 312.7267993 |
| 644.1010927 | 84.55993425 |
| 655.4374068 | 358.1991539 |
| 719.5778962 | 178.1754498 |
| 759.2255662 | 250.2589341 |
| 883.7443645 | 26.3450723 |
| 935.8756609 | 131.4753467 |
| 964.5537569 | 309.1573473 |
| 1001.544653 | 204.4847256 |
| 1109.337326 | 71.40197912 |
| 1165.076192 | 166.8240659 |
| 1173.018943 | 346.932216 |
| 1192.049242 | 255.5296653 |
| 1250.993603 | 113.3983338 |
| 1252.645007 | 198.036268 |
| 1255.216795 | 22.46042154 |
| 1299.899795 | 304.4751002 |
| 1331.279187 | 142.7238208 |
| 1338.3037 | 60.03285078 |
| 1350.249082 | 341.3462322 |
| 1367.034769 | 242.2954933 |
| -1 | |

1.33

| | |
|---|---|
| 251.3247643 | 320.9860399 |
| 265.7891414 | 52.04964138 |
| 274.2139882 | 227.3171352 |
| 294.1866052 | 140.8869546 |
| 451.3604625 | 274.2963231 |
| 567.6791646 | 181.6085305 |
| 567.8930934 | 101.4344101 |
| 570.3109376 | 357.5403189 |
| 763.9343403 | 44.9158703 |
| 855.4717672 | 231.9981719 |
| 900.4779507 | 307.1521215 |
| 917.0773391 | 147.8422855 |
| 1089.70625 | 18.33302093 |
| 1137.614959 | 82.00089364 |
| 1197.008235 | 266.8776953 |
| 1248.888957 | 196.3089356 |
| 1422.782381 | 134.701726 |
| 1445.801794 | 327.428006 |
| 1551.871387 | 229.5606565 |
| 1563.575182 | 45.03613444 |
| 1684.064606 | 281.838515 |
| 1685.478988 | 169.3029807 |
| 1695.221423 | 88.69017644 |
| 1738.539297 | 351.5000477 |
| 1769.254801 | 125.0985806 |
| 1794.352139 | 318.0305097 |
| 1798.700724 | 51.42782711 |
| 1806.317188 | 204.0992606 |
| 1813.723946 | 251.7989176 |

-1

1.35

| | |
|---|---|
| 172.5602644 | 0.05724691675 |
| 173.9298897 | 102.6487236 |
| 179.2475725 | 205.4676757 |
| 193.3293464 | 287.0201653 |
| 294.5415528 | 52.74667512 |
| 306.9333605 | 152.7786514 |
| 387.8606396 | 345.6522637 |
| 391.5945847 | 227.6802908 |
| 447.8922432 | 96.20632993 |
| 502.2365765 | 289.4437331 |

| | |
|---|---|
| 550.3122444 | 36.78618564 |
| 556.0270009 | 172.9178764 |
| 666.8030208 | 236.5396588 |
| 671.9036689 | 348.4027301 |
| 672.920274 | 111.0719685 |
| 740.7600287 | 294.5785105 |
| 764.2657755 | 54.68886542 |
| 788.8711368 | 174.5176084 |
| 826.0589038 | 356.8704243 |
| 826.0819692 | 120.0465905 |
| 843.6993228 | 236.9988792 |
| -1 | |
| 1.36 | |
| 152.3377996 | 59.37758268 |
| 153.2453541 | 151.1455959 |
| 154.9939927 | 242.6966182 |
| 159.2063639 | 336.928507 |
| 276.1220004 | 291.0511242 |
| 279.4538773 | 102.5690769 |
| 298.6722665 | 188.8063693 |
| 302.6168313 | 24.73223745 |
| 391.2369962 | 239.7748441 |
| 455.6613055 | 141.9738864 |
| 462.7841994 | 332.1230819 |
| 480.034464 | 62.72859644 |
| 554.7626014 | 277.6762916 |
| 575.6264273 | 198.7470304 |
| 589.3638285 | 15.71905544 |
| 615.5539883 | 114.7619583 |
| 629.3882776 | 320.8144737 |
| 669.8751624 | 227.4595853 |
| 679.4048741 | 63.41511924 |
| 703.4912815 | 158.0173438 |
| -1 | |
| 1.38 | |
| 138.3564989 | 190.1510996 |
| 142.0882552 | 1.269125143 |
| 144.1065138 | 276.224269 |
| 158.5263193 | 99.2681918 |
| 274.8178313 | 143.4570319 |
| 276.738884 | 49.86864451 |
| 288.0321912 | 233.2706463 |

| | |
|---|---|
| 295.4852551 | 319.3075641 |
| 406.5276505 | 97.27267648 |
| 412.8175164 | 15.10634238 |
| 421.850415 | 181.9984768 |
| 444.1817476 | 268.7525408 |
| 485.167619 | 321.4379626 |
| 505.9681769 | 69.19644118 |
| 507.1651332 | 136.0387199 |
| 518.0665277 | 222.7888053 |
| 523.1838849 | 14.08460958 |
| 556.3019142 | 174.256799 |
| 556.7888623 | 283.7612985 |
| 564.2185861 | 105.5687167 |
| -1 | |
| 1.39.2 | |
| 219.7091907 | 311.7723844 |
| 220.5434215 | 13.899233 |
| 251.7814185 | 93.31633562 |
| 259.641742 | 173.1512647 |
| 259.6880015 | 235.8932513 |
| 445.8201611 | 150.0260379 |
| 452.0186524 | 275.6397188 |
| 487.2859561 | 346.5615167 |
| 490.111318 | 59.64614201 |
| 506.6039387 | 206.672145 |
| 669.7358042 | 113.0357893 |
| 703.4881581 | 29.85513948 |
| 727.0040191 | 319.2060855 |
| 762.2158688 | 173.5979791 |
| 764.4204496 | 249.8706352 |
| 907.4044382 | 61.16838441 |
| 948.298192 | 212.5758642 |
| 956.2265377 | 1.521730205 |
| 976.6520523 | 132.2962206 |
| 989.3606219 | 302.1615874 |
| 1111.58899 | 255.3079136 |
| 1113.16703 | 48.06181039 |
| 1114.248782 | 183.9138944 |
| 1122.254784 | 343.4215059 |
| 1171.553783 | 118.7560553 |
| 1188.507649 | 299.7081768 |
| 1211.678469 | 165.6722315 |

| | |
|---|---|
| 1212.359195 | 28.32210959 |
| -1 | |
| 1.37 | |
| 165.4617286 | 183.9825573 |
| 174.1831397 | 97.24216055 |
| 188.737198 | 305.30224 |
| 209.513012 | 5.919055786 |
| 288.1478049 | 233.9849824 |
| 327.0000887 | 335.6284346 |
| 330.4744843 | 149.9158114 |
| 380.3738631 | 53.14928768 |
| 433.548829 | 279.1196299 |
| 462.8611778 | 190.1612919 |
| 466.3059016 | 4.90237167 |
| 533.1774617 | 118.0734178 |
| 601.0779316 | 339.3085375 |
| 613.8497163 | 215.462412 |
| 639.0261528 | 38.1658593 |
| 680.7615755 | 74.23692082 |
| 727.5729569 | 292.4277864 |
| 760.663763 | 1.483292931 |
| 778.7844053 | 111.209809 |
| 780.8498668 | 195.9038632 |
| 806.7495199 | 342.9186331 |
| 824.8011011 | 232.2011371 |
| 848.5346206 | 34.28323721 |
| 854.1904945 | 153.0577883 |
| -1 | |
| 1.40 | |
| 166.0851007 | 180.6070535 |
| 169.7070203 | 279.9142173 |
| 178.2396428 | 32.54740433 |
| 223.8027838 | 105.8781306 |
| 290.7567656 | 232.5394895 |
| 306.9106629 | 331.8307215 |
| 384.1296688 | 164.5127576 |
| 390.8204399 | 54.97576932 |
| 433.4634015 | 277.4555932 |
| 499.9848673 | 108.0438231 |
| 537.2743916 | 215.5496457 |
| 555.6792041 | 350.6024249 |
| 661.1681809 | 292.4170867 |

| | |
|---|---|
| 666.2351801 | 52.0358525 |
| 667.6585543 | 162.3506764 |
| 738.4928387 | 111.6571138 |
| 757.7589858 | 237.9792994 |
| 773.862175 | 356.1194152 |
| 818.9708557 | 176.5296897 |
| 820.2917233 | 300.8314759 |
| 842.5054891 | 58.57799852 |
| -1 | |
| 1.41 | |
| 215.1993061 | 23.8596982 |
| 251.2142619 | 96.08250951 |
| 251.5845176 | 163.732628 |
| 254.0216407 | 235.1949807 |
| 258.4660936 | 308.9365248 |
| 453.2966248 | 352.6657308 |
| 463.2597067 | 57.20291155 |
| 518.0230194 | 126.5470342 |
| 545.2680762 | 281.244683 |
| 545.9953837 | 201.8313777 |
| 762.0686466 | 329.7027094 |
| 792.1629777 | 34.44788881 |
| 837.0211604 | 162.5609499 |
| 837.4393581 | 98.20769237 |
| 840.7194297 | 248.2926578 |
| 1050.39829 | 4.984467219 |
| 1057.545584 | 291.5217255 |
| 1119.10821 | 147.8098053 |
| 1139.722618 | 77.01741781 |
| 1157.285979 | 214.4475005 |
| 1226.42107 | 332.8762506 |
| 1284.073749 | 266.7224629 |
| 1291.250913 | 28.61787014 |
| 1324.554536 | 112.7479271 |
| 1346.651091 | 179.1107669 |
| 1370.452847 | 301.5480952 |
| 1378.090389 | 230.8756169 |
| 1401.508946 | 71.55171318 |
| -1 | |
| 1.43 | |
| 132.7945379 | 145.3052243 |
| 135.451834 | 321.3257822 |

| | |
|---|---|
| 152.1230422 | 57.84380882 |
| 153.4618531 | 224.325443 |
| 277.9552335 | 280.6600025 |
| 279.6764147 | 1.715103991 |
| 289.6524048 | 105.3934713 |
| 331.7636507 | 193.9923328 |
| 367.9335216 | 55.55910713 |
| 397.0681786 | 318.5652265 |
| 401.0245878 | 140.7009839 |
| 406.7667994 | 243.6222472 |
| 423.8735569 | 195.9772129 |
| 445.5562138 | 84.83458343 |
| 446.4681648 | 19.55417063 |
| 453.5210108 | 279.5177218 |
| -1 | |
| 1.45 | |
| 206.3561965 | 94.74792896 |
| 224.0504523 | 182.3579732 |
| 226.8863998 | 354.5325192 |
| 252.3547668 | 258.1628639 |
| 454.9490704 | 54.53634267 |
| 482.2909475 | 222.3761461 |
| 526.6499707 | 300.9509374 |
| 574.6957514 | 136.4824876 |
| 728.4018627 | 179.2232662 |
| 744.6368156 | 356.6628209 |
| 814.6273047 | 67.97919531 |
| 816.436277 | 253.7299263 |
| 1081.662139 | 306.3866452 |
| 1098.093299 | 127.5834836 |
| 1131.490015 | 205.1647728 |
| 1164.588774 | 24.41940339 |
| 1192.293076 | 78.3739355 |
| 1339.988133 | 343.3804416 |
| 1348.440318 | 262.7056512 |
| 1354.517623 | 164.7422907 |
| 1364.332423 | 78.67754714 |
| 1395.701049 | 295.6924222 |
| 1397.615098 | 208.5038918 |
| 1429.552449 | 120.9525939 |
| 1473.4679 | 28.84218076 |
| 1507.847806 | 338.2292605 |

−1
1.46

| | |
|---|---|
| 185.1225453 | 196.2550911 |
| 198.7063409 | 6.940368937 |
| 209.0946428 | 121.3400757 |
| 213.3744293 | 279.7688398 |
| 321.8630074 | 66.6480738 |
| 407.5944818 | 156.1070783 |
| 421.9841199 | 321.4827206 |
| 460.5071501 | 241.428314 |
| 492.888276 | 18.31948404 |
| 547.4795483 | 109.2209233 |
| 695.1251181 | 196.2550911 |
| 698.7584379 | 62.0420867 |
| 729.6945262 | 280.7862753 |
| 807.9912453 | 155.1907476 |
| 814.3340199 | 345.7954411 |
| 898.735896 | 103.8609531 |
| 931.1478486 | 34.74337655 |
| 933.8531528 | 247.6689477 |
| 1007.401255 | 303.8219765 |
| 1020.776148 | 191.3959413 |
| 1036.118404 | 76.68154693 |
| 1045.125361 | 355.5443014 |
| 1062.894899 | 134.7122097 |
| 1082.382319 | 255.4145007 |

−1
1.47

| | |
|---|---|
| 235.6974957 | 63.6712239 |
| 248.2356483 | 286.8247778 |
| 248.3460602 | 149.3199625 |
| 277.1112862 | 353.1906392 |
| 291.5383548 | 211.9850854 |
| 461.73904 | 118.3986289 |
| 547.4189948 | 177.5977877 |
| 548.1552962 | 33.18884384 |
| 564.8926999 | 319.803247 |
| 646.2925775 | 247.1596455 |
| 879.2636138 | 29.47934053 |
| 882.0781948 | 123.3409305 |
| 939.6329624 | 334.2937608 |
| 985.8811641 | 210.2441038 |

| | |
|---|---|
| 1082.474412 | 278.1014272 |
| 1206.217687 | 134.5638094 |
| -1 | |
| 1.48 | |
| 224.1134888 | 245.1500384 |
| 226.0048895 | 314.4837389 |
| 228.9370717 | 18.00401464 |
| 237.6436529 | 162.6290128 |
| 253.3587146 | 101.2221055 |
| 426.3797068 | 203.4775597 |
| 428.0338917 | 346.0304183 |
| 440.5930472 | 271.1902355 |
| 490.6063752 | 63.65616889 |
| 522.1703952 | 129.1071186 |
| 724.3165163 | 232.8610522 |
| 726.6652004 | 319.635313 |
| 810.223616 | 23.04931237 |
| 811.1160682 | 97.82672952 |
| 816.7451245 | 172.5643025 |
| 983.6586961 | 279.2709838 |
| 1033.092417 | 127.7988414 |
| 1053.735527 | 59.27275276 |
| 1060.105699 | 335.6881268 |
| 1088.53218 | 205.3905667 |
| 1165.430107 | 139.1617341 |
| 1167.14246 | 262.2733929 |
| 1181.221247 | 6.405789608 |
| 1202.529449 | 73.44007895 |
| 1236.535606 | 206.3583458 |
| -1 | |
| 1.52 | |
| 210.0073722 | 174.9779572 |
| 242.0818044 | 105.3724858 |
| 245.52731 | 259.0798381 |
| 284.984739 | 338.9618307 |
| 291.821286 | 38.17200803 |
| 397.9704256 | 217.706633 |
| 439.1641443 | 126.0315951 |
| 445.6519263 | 60.81465365 |
| 454.3770858 | 303.1559416 |
| 574.0830748 | 3.97819648 |
| 585.3156901 | 166.8510529 |

| | |
|---|---|
| 621.9125421 | 250.3962391 |
| 699.1276314 | 86.97825179 |
| 758.0228975 | 199.4411409 |
| 787.3236724 | 316.7998906 |
| 849.9461827 | 28.64203128 |
| 858.5845107 | 138.5359037 |
| 926.9272047 | 231.6280917 |
| 1016.049628 | 181.4835734 |
| 1017.556337 | 86.29175932 |
| 1037.840403 | 348.5980284 |
| 1076.597108 | 280.9381809 |
| 1110.270435 | 213.83546 |
| 1131.125618 | 144.5829542 |
| 1176.114391 | 37.90887201 |
| 1211.305719 | 249.8533516 |
| 1218.494392 | 178.9781988 |
| 1237.020104 | 324.678571 |
| 1255.240657 | 80.80955084 |
| 1342.753356 | 12.43426813 |
| -1 | |
| 1.54 | |
| 106.7049988 | 238.4073085 |
| 130.6037561 | 118.8207637 |
| 135.0739572 | 351.3097119 |
| 153.15467 | 182.6994188 |
| 184.6902545 | 36.09479657 |
| 188.1576546 | 304.9424858 |
| 222.601473 | 226.0522406 |
| 229.6716519 | 130.5222267 |
| 248.5124381 | 175.5875246 |
| 258.3798045 | 351.2734811 |
| 267.2873713 | 51.33631673 |
| 270.2090986 | 296.6937197 |
| -1 | |
| 1.55 | |
| 102.8300414 | 249.9690752 |
| 117.8117018 | 70.28249054 |
| 123.3893571 | 338.1003288 |
| 133.0210319 | 167.1035167 |
| 185.5238449 | 25.48149846 |
| 194.0979967 | 294.3655544 |
| 197.1399974 | 224.0567791 |

| | |
|---|---|
| 197.4409728 | 124.5867822 |
| 227.3434765 | 340.6162624 |
| 233.6078081 | 170.4667489 |
| 235.5427845 | 79.48162401 |
| 237.0292328 | 247.1650493 |
| 258.0119671 | 21.88753013 |
| 262.2803125 | 155.5885785 |
| 263.5378882 | 310.1250832 |
| -1 | |
| 1.56 | |
| 94.40994601 | 295.1307709 |
| 99.48867167 | 207.8114795 |
| 105.8846647 | 30.73824613 |
| 116.1942131 | 118.6191147 |
| 176.1651771 | 86.4726005 |
| 178.0471717 | 261.6534373 |
| 180.3370394 | 339.819795 |
| 181.3076665 | 220.8155388 |
| 182.051644 | 140.0661853 |
| 194.7474368 | 18.3208586 |
| 195.741911 | 297.0850213 |
| -1 | |
| 1.58 | |
| 245.0975859 | 203.8922958 |
| 267.1670412 | 6.199703715 |
| 277.5214353 | 282.0380357 |
| 283.2757353 | 137.3633158 |
| 297.0612984 | 82.68654439 |
| 516.1416407 | 244.5166252 |
| 534.1626362 | 161.8837937 |
| 546.934151 | 338.5896939 |
| 563.1523853 | 43.12325982 |
| 671.5566818 | 288.5704242 |
| 702.9315419 | 109.2366958 |
| 739.2885678 | 200.0961283 |
| 827.6757902 | 15.00590561 |
| 899.0982693 | 244.7231061 |
| 915.7785185 | 152.4622244 |
| 973.9667532 | 73.70866739 |
| 982.0991202 | 325.9397696 |
| 1086.442008 | 193.372971 |
| 1101.221034 | 124.0348621 |

| | |
|---|---|
| 1118.071807 | 32.03047391 |
| 1121.296865 | 283.2492323 |
| 1143.875857 | 235.5945236 |
| 1186.610292 | 159.9215269 |
| 1209.272471 | 87.59674258 |
| 1212.287571 | 351.5094777 |
| 1213.539385 | 308.8812635 |
| 1247.460501 | 44.16036475 |
| -1 | |
| 1.59 | |
| 115.6947352 | 253.0775521 |
| 117.9881635 | 139.2812085 |
| 134.2565378 | 343.617188 |
| 137.6439744 | 65.94343679 |
| 190.8923896 | 184.8031959 |
| 210.0260063 | 100.6380757 |
| 218.9605331 | 302.5029043 |
| 243.6341223 | 19.73414836 |
| 245.3782887 | 141.8990551 |
| 258.9131143 | 247.887999 |
| 270.2189046 | 85.66720678 |
| 280.4246977 | 185.4896325 |
| 285.6101611 | 322.0666399 |
| 294.4986758 | 18.28700401 |
| -1 | |
| 1.60.3 | |
| 148.3250835 | 176.6068526 |
| 158.8574206 | 327.8332207 |
| 162.8498988 | 81.24047917 |
| 172.4571121 | 18.58332178 |
| 173.5372346 | 110.9864479 |
| 178.5271019 | 242.853152 |
| 303.151065 | 145.2941951 |
| 362.2284235 | 35.48106755 |
| 377.5932888 | 216.0818682 |
| 406.4818651 | 335.8514788 |
| 517.953822 | 271.6080214 |
| 538.4043736 | 95.67205131 |
| 617.0143241 | 171.7193837 |
| 722.940081 | 5.667635549 |
| 784.6415177 | 55.22636105 |
| 852.2849362 | 215.8511597 |

| | |
|---|---|
| 920.6954923 | 138.710194 |
| 933.4847733 | 317.3012697 |
| 1099.003916 | 5.671302165 |
| 1106.409565 | 185.8606577 |
| 1114.294438 | 84.99976393 |
| 1154.178622 | 264.1352824 |
| 1237.043809 | 137.1487111 |
| 1251.499929 | 42.68545872 |
| 1269.952447 | 224.2077972 |
| 1278.137248 | 310.3590586 |
| −1 | |
| 1.62.1 | |
| 120.3945084 | 317.8391762 |
| 121.1577448 | 241.1189896 |
| 124.1378785 | 54.79015031 |
| 124.5678868 | 161.583945 |
| 181.826082 | 110.1245202 |
| 183.1252057 | 2.010998083 |
| 185.30722 | 204.4376358 |
| 195.5229301 | 279.0594231 |
| 220.1434609 | 58.03378761 |
| 235.6087842 | 145.349903 |
| 249.0529534 | 338.9705215 |
| 250.1591478 | 238.550829 |
| 270.7538571 | 78.21796641 |
| 275.8154148 | 175.337895 |
| 281.5561713 | 293.1459415 |
| −1 | |
| 1.62.3 | |
| 149.0798159 | 249.3021939 |
| 161.8357304 | 33.56245188 |
| 162.7158074 | 132.7476828 |
| 164.9779241 | 308.9226226 |
| 216.7642756 | 202.9897148 |
| 226.6602252 | 82.34113275 |
| 262.9050413 | 343.8839081 |
| 297.1621473 | 271.1702229 |
| 299.7161108 | 149.5464542 |
| 349.8765143 | 103.7824002 |
| 374.8996553 | 42.54475895 |
| 416.6687686 | 214.4991515 |
| 421.07406 | 314.4744692 |

| | |
|---|---|
| 474.3134393 | 138.1235092 |
| 486.3009311 | 35.1191257 |
| 488.2742596 | 263.5629765 |
| 501.1260327 | 94.25036034 |
| 511.707477 | 332.2100742 |
| 532.112551 | 184.907865 |
| 545.0161983 | 253.4172612 |
| 560.5959034 | 33.31499121 |
| 568.6066862 | 121.2421741 |
| -1 | |
| 1.63 | |
| 213.4874263 | 206.9167714 |
| 231.891027 | 340.8436209 |
| 238.1387419 | 128.3582698 |
| 238.8712886 | 268.2658489 |
| 243.4440008 | 57.40207042 |
| 407.8065875 | 11.66527118 |
| 410.5323639 | 168.3722891 |
| 422.6163208 | 300.052766 |
| 439.4645482 | 242.5963762 |
| 447.8995312 | 82.98886508 |
| 664.7655806 | 204.4849282 |
| 672.05187 | 274.0095121 |
| 682.6561578 | 343.5797334 |
| 704.2549231 | 127.9549425 |
| 742.4754392 | 44.54757554 |
| 865.6798841 | 246.0309564 |
| 874.8816459 | 77.95139336 |
| 883.971538 | 164.1563201 |
| 921.8650274 | 309.3464273 |
| 970.3920137 | 26.55845676 |
| 992.5824125 | 205.9448949 |
| 1017.977533 | 108.376296 |
| 1028.06022 | 271.3736864 |
| 1051.968697 | 343.9314163 |
| 1052.709166 | 66.66689667 |
| 1084.520877 | 160.6745898 |
| -1 | |
| 1.66 | |
| 120.023309 | 338.8639602 |
| 120.2240778 | 202.4708519 |
| 132.2349276 | 72.57743984 |

| | |
|---|---|
| 181.077544 | 8.135694956 |
| 190.5331516 | 267.1878704 |
| 195.8571008 | 151.3869496 |
| 210.2904453 | 328.0410191 |
| 212.1210194 | 76.8301428 |
| 220.4426701 | 195.6621147 |
| 239.1526259 | 10.85378576 |
| 243.191788 | 272.6200919 |
| 255.1069014 | 149.3541191 |
| -1 | |
| 1.68 | |
| 117.253427 | 31.49108077 |
| 126.6130203 | 313.7704397 |
| 127.7786955 | 127.4067015 |
| 131.9485215 | 204.1353968 |
| 169.3715666 | 356.0278268 |
| 170.2668537 | 261.0430438 |
| 172.5094151 | 73.87263748 |
| 176.2330636 | 164.1202722 |
| 237.2934765 | 209.6696417 |
| 238.6183525 | 316.3703156 |
| 247.6263375 | 113.2322345 |
| 252.7082197 | 27.22833049 |
| 292.7141371 | 257.3977031 |
| 301.5970187 | 349.4080969 |
| 305.6372972 | 169.4313826 |
| 317.2823342 | 73.98594106 |
| -1 | |
| 1.69 | |
| 121.2945701 | 100.2767554 |
| 123.2210063 | 11.73032764 |
| 130.3437014 | 189.2419121 |
| 133.1718027 | 288.019427 |
| 220.8619454 | 60.51111392 |
| 227.5867624 | 332.8590977 |
| 235.9524603 | 236.0628455 |
| 244.8051569 | 149.4161834 |
| 298.4832749 | 191.7664805 |
| 300.490569 | 288.6811678 |
| 314.9235031 | 102.5825408 |
| 322.6706029 | 8.199981271 |
| 353.3161646 | 240.0197185 |

| | |
|---|---|
| 367.0959835 | 353.3358083 |
| 367.2361641 | 63.61528023 |
| 369.1470241 | 155.6225138 |
| -1 | |
| 1.70 | |
| 197.5581908 | 21.04401567 |
| 199.735828 | 165.5976493 |
| 208.7599127 | 223.9760568 |
| 210.5289933 | 307.6411689 |
| 215.2065824 | 95.59965205 |
| 368.2893395 | 344.139299 |
| 372.025307 | 53.3935214 |
| 388.1271342 | 195.4922817 |
| 394.8917837 | 136.8755544 |
| 409.9658747 | 268.9849863 |
| 609.4637097 | 19.39380124 |
| 610.5728491 | 319.4692329 |
| 610.5856639 | 95.43964771 |
| 632.7974535 | 219.1770025 |
| 684.3690088 | 161.0655617 |
| 746.9605764 | 268.9850267 |
| 805.3481605 | 43.04491555 |
| 807.6425781 | 343.3906201 |
| 857.4432396 | 126.4113043 |
| 879.9362044 | 194.691516 |
| 893.3611987 | 9.258753068 |
| 894.4919328 | 272.6698862 |
| 924.9342691 | 80.75575777 |
| 931.432099 | 220.0677351 |
| 946.9842551 | 158.7302613 |
| -1 | |
| 1.74 | |
| 138.7137522 | 189.8363557 |
| 139.8886791 | 96.40212317 |
| 141.7554374 | 5.297719408 |
| 149.6370875 | 276.5993079 |
| 229.886423 | 322.0534573 |
| 237.4457504 | 59.23814628 |
| 255.0137725 | 227.9038854 |
| 284.455769 | 140.7482032 |
| 305.9395085 | 17.16810772 |
| 320.252745 | 275.1954394 |

| | |
|---|---|
| 347.4303628 | 92.54071648 |
| 367.6738633 | 185.058913 |
| 381.0238213 | 323.6342409 |
| 411.1840652 | 46.24186425 |
| 430.1871333 | 235.2796513 |
| 448.5481748 | 5.506306989 |
| 461.7242493 | 288.4041207 |
| 463.1718134 | 132.9275816 |
| 491.172759 | 80.73555821 |
| 497.595358 | 183.5278817 |
| -1 | |
| 1.75 | |
| 137.5061585 | 94.71170691 |
| 140.7036397 | 209.0221611 |
| 143.7698264 | 328.0501855 |
| 201.1666236 | 38.08831273 |
| 216.1563869 | 140.3534626 |
| 248.0549141 | 276.3839149 |
| 267.9228375 | 82.2561569 |
| 311.1871927 | 346.1166234 |
| 318.1580265 | 212.0742087 |
| 373.9793463 | 157.9242694 |
| 389.2521847 | 37.3380447 |
| 397.2499215 | 281.2371332 |
| 429.7529834 | 97.67190526 |
| 463.5616653 | 216.5587889 |
| 467.1362594 | 339.6469177 |
| 494.4715693 | 51.50301393 |
| 515.3920244 | 154.6367192 |
| 521.8166305 | 103.911694 |
| 530.0660955 | 272.8508183 |
| 536.688016 | 355.2485184 |
| 567.4611816 | 219.8777961 |
| -1 | |
| 1.81 | |
| 130.4123754 | 359.5330282 |
| 137.6980171 | 245.2273449 |
| 142.1194021 | 136.2919649 |
| 153.326001 | 58.5350486 |
| 222.4432233 | 303.7225052 |
| 236.9448386 | 183.3988845 |
| 261.7481354 | 80.73810004 |

| | |
|---|---|
| 297.5432868 | 7.719664269 |
| 300.9617394 | 245.5150334 |
| 334.3362325 | 140.7153806 |
| 389.9624604 | 51.73312114 |
| 391.6607554 | 195.5975684 |
| 391.9251737 | 327.1798721 |
| 439.6656161 | 6.920365038 |
| 442.2775464 | 96.86652602 |
| 444.3060079 | 273.7786572 |
| 463.6881907 | 158.2530834 |
| 491.4191289 | 62.6875619 |
| 492.3542879 | 319.2046476 |
| 505.1362467 | 222.6298063 |
| -1 | |
| 1.81.2 | |
| 136.8377276 | 197.5670512 |
| 141.3520814 | 343.8617444 |
| 146.5574335 | 86.89466163 |
| 181.131062 | 260.5232662 |
| 210.7025274 | 21.15244951 |
| 211.8148348 | 140.1138668 |
| 262.3184594 | 197.4899026 |
| 269.485221 | 331.8523098 |
| 293.8171472 | 85.86325669 |
| 342.9359131 | 262.9033511 |
| 345.8710304 | 142.5782817 |
| 367.6036392 | 18.84255537 |
| 400.6705325 | 201.101335 |
| 416.2028681 | 82.81050932 |
| 433.9176577 | 323.3902651 |
| 464.7189332 | 140.0030552 |
| 474.7395961 | 19.76749941 |
| 485.5270329 | 264.1020026 |
| 505.8642987 | 75.90413632 |
| 520.6937316 | 196.6827991 |
| 547.3827189 | 328.1245759 |
| -1 | |
